# Extracellular matrix supports excitation-inhibition balance in neuronal networks by stabilizing inhibitory synapses

**DOI:** 10.1101/2020.07.13.200113

**Authors:** Egor Dzyubenko, Michael Fleischer, Daniel Manrique-Castano, Mina Borbor, Christoph Kleinschnitz, Andreas Faissner, Dirk M Hermann

**Affiliations:** Department of Neurology, University Hospital Essen, Hufelandstraße 55, D-45122 Essen, Germany; Department of Cell Morphology and Molecular Neurobiology, Faculty of Biology and Biotechnology, Ruhr University Bochum, Universitätsstraße 150, D-44801 Bochum, Germany

## Abstract

Maintaining the balance between excitation and inhibition is essential for the appropriate control of neuronal network activity. Sustained excitation-inhibition (E-I) balance relies on the orchestrated adjustment of synaptic strength, neuronal activity and network circuitry. While growing evidence indicates that extracellular matrix (ECM) of the brain is a crucial regulator of neuronal excitability and synaptic plasticity, it remains unclear whether and how ECM contributes to neuronal circuit stability. Here we demonstrate that the integrity of ECM supports the maintenance of E-I balance by retaining inhibitory connectivity. Depletion of ECM in mature neuronal networks preferentially decreases the density of inhibitory synapses and the size of individual inhibitory postsynaptic scaffolds. After ECM depletion, inhibitory synapse strength homeostatically increases via the reduction of presynaptic GABA_B_ receptors. However, the inhibitory connectivity reduces to an extent that inhibitory synapse scaling is no longer efficient in controlling neuronal network activity. Our results indicate that the brain ECM preserves the balanced network state by stabilizing inhibitory synapses.

**Significance statement:** The question how the brain’s extracellular matrix (ECM) controls neuronal plasticity and network activity is key for an appropriate understanding of brain functioning. In this study, we demonstrate that ECM depletion much more strongly affects the integrity of inhibitory than excitatory synapses in vitro and in vivo. We revealed that by retaining inhibitory connectivity, ECM ensures the efficiency of inhibitory control over neuronal network activity. Our work significantly expands our current state of knowledge about the mechanisms of neuronal network activity regulation. Our findings are similarly relevant for researchers working on the physiological regulation of neuronal plasticity in vitro and in vivo and for researchers studying the remodeling of neuronal networks upon brain injury, where prominent ECM alterations occur.

## Introduction

Neuronal network activity is regulated through a dynamic balance of excitation and inhibition (Haider et al., 2006) that requires coordinated plasticity of excitatory and inhibitory synapses (Bhatia et al., 2019; Trapp et al., 2018). Over several years, experimental studies have gathered solid evidence on the plasticity of excitatory synapses, establishing the modulation of presynaptic neurotransmitter release and postsynaptic responsiveness to glutamate as key neural correlates for memory and learning (Ho et al., 2011). On the network level, scaling of inhibitory synapses is essential for homeostatic mechanisms maximizing information processing capacity in neuronal networks (Ma et al., 2019). Computational modelling suggests that patterns of neuronal network activity are primarily determined by inhibitory connectivity (Mongillo et al., 2018). Yet, the limited knowledge about inhibitory synapses (for review see Gandolfi et al., 2020) limits our understanding how inhibitory plasticity and connectivity influence neuronal activity at the integrative network level.

The brain extracellular matrix (ECM) is a multicomponent macromolecular meshwork containing chondroitin sulfate carrying proteoglycans (CSPGs), which anchors to the neuronal surface via hyaluronic acid synthesizing enzymes (Krishnaswamy et al., 2019; Roll and Faissner, 2014). Neuronal activity induces the consolidation of ECM molecules (Dityatev et al., 2007) forming densely packed lattice-shaped layers around a subpopulation of neurons. These coatings, termed perineuronal nets (PNNs), support the fast spiking properties of interneurons (Chu et al., 2018), regulate learning and help to retain acquired memory (Carulli et al., 2020). In adulthood, CSPGs of PNNs and interstitial ECM restrict neuronal plasticity (Carulli et al., 2006; Carulli et al., 2007; Dzyubenko et al., 2016a; Pizzorusso et al., 2002).

Enzymatic digestion of ECM increases lateral mobility of excitatory glutamate receptors, alters short-term plasticity of excitatory synapses (Frischknecht et al., 2009) and enhances neuronal network activity (Bikbaev et al., 2015). Thus, ECM integrity is essential for sustained synaptic signaling and neuronal circuit stability. Upon brain injury induced by ischemic stroke, the ECM is rapidly degraded within a few hours (Härtig et al., 2017), and it remains partly decomposed after one week in lesion-surrounding brain areas (Dzyubenko et al., 2018), in which neuronal network activity is compromised (Clarkson et al., 2010; Lake et al., 2017). Whether ECM depletion contributes to the altered neuronal network activity after brain injury and how it influences synaptic plasticity remains to be identified. The role of inhibitory synapses in controlling neuronal network activity after ECM depletion was unknown.

Here we investigated the role of ECM for stabilizing excitatory and inhibitory synapses and balancing neuronal network activity. We enzymatically degraded hyaluronic acid and CSPGs in the extracellular space and combined structural and functional readouts for studying neuronal connectivity and activity at the synapse and network level. By using data-driven computer simulations, we explored the influence of synaptic strength and connectivity changes on the activity of neuronal networks.

## Results

### ECM depletion reduces inhibitory synapse density *in vitro*

Measuring synapse density changes provides an indirect but straightforward estimate of neuronal network connectivity alterations (Dzyubenko et al., 2017). Thus, we first quantified the density of structurally complete synapses in mature networks of primary murine neurons. The networks consisted of principal excitatory neurons and inhibitory interneurons in the ratio 2:1 and were fully mature after 21 days of cultivation, indicated by the appearance of PNNs, which are condensed ECM layers, in neuronal cultures (Figures S1 and S2). The density of both GABAergic and glutamatergic synapses was quantified using GABA as marker of inhibitory perikarya, allowing for the gross estimation of network connectivity. The co-labelling of vesicular glutamate transporter type 1 (VGLUT1) and postsynaptic density protein 95 (PSD95) indicated the excitatory inputs to excitatory and inhibitory neurons, while the co-labelling of vesicular GABA transporter (VGAT) and gephyrin signified inhibitory inputs to excitatory and inhibitory neurons (Figure 1A, B).

**Figure 1.**
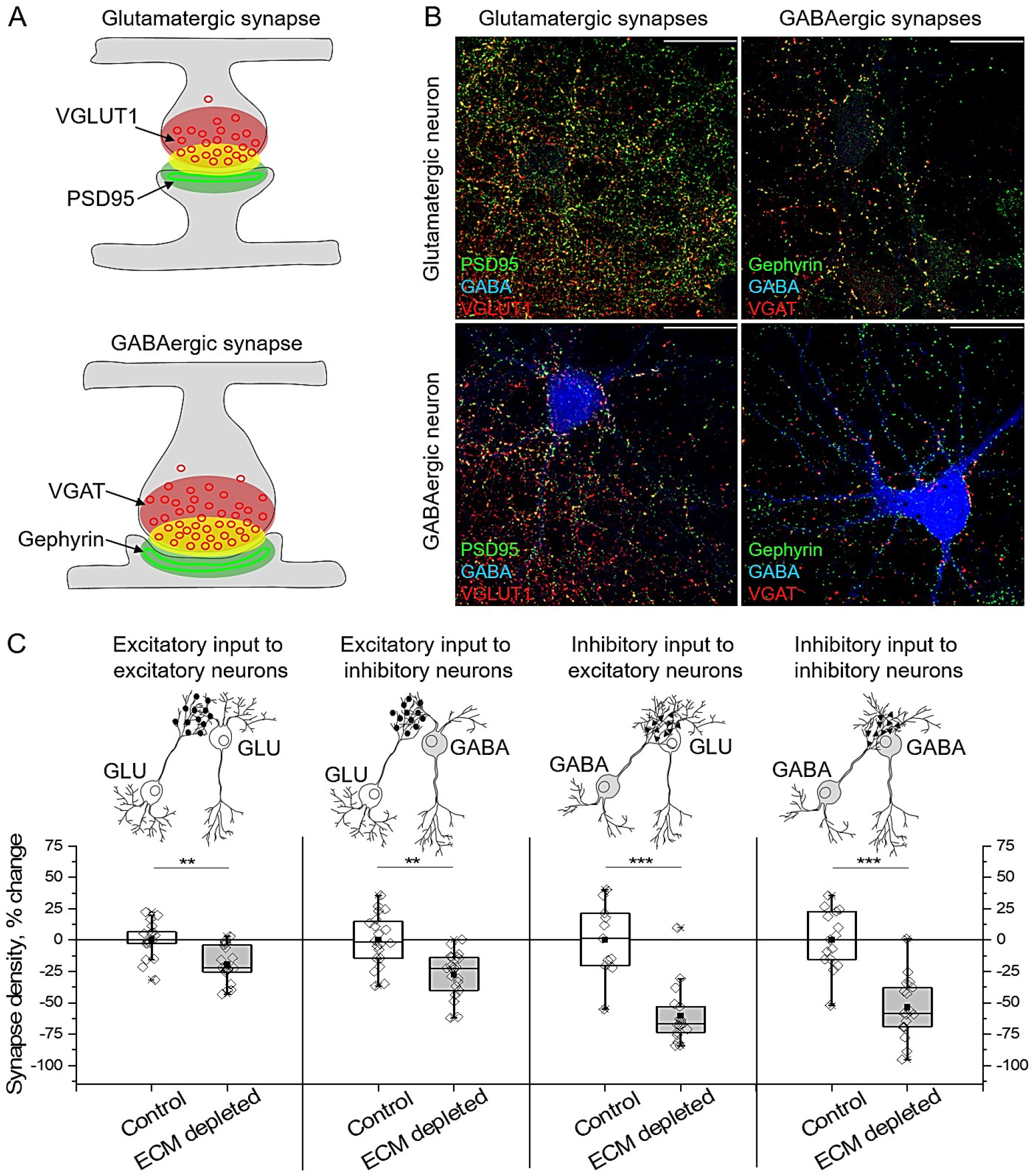
Excitatory and inhibitory synapse densities decrease after ECM depletion *in vitro.* (A) Overlapping immunolabelling of presynaptic (red) and postsynaptic (green) markers was used to detect structurally complete synapses (yellow). (B) The density of glutamatergic (PSD95-VGLUT1) and GABAergic (gephyrin-VGAT) synapses was measured with reference to GABA immunoreactivity. Representative micrographs are shown. Scale bars, 30 µm. (C) Synapse density changes were calculated as differences with mean values of corresponding control experiments. Data are shown for each neuron examined (n≥20 neurons per condition, results obtained from 5 independent experiments). GLU, glutamate. Data are medians (lines inside boxes)/ means (filled squares inside boxes) ± IQR (boxes) with 10/ 90% ranks as whiskers. Open diamonds are data points. The asterisks indicate significant differences with control, based on Kruskal-Wallis tests (***p<0.001, **p<0.01).

The density of both glutamatergic and GABAergic synapses decreased after enzymatic ECM depletion (500 mU/ml chondroitinase ABC [ChABC], 16 hours). Compared with control, ECM depletion reduced excitatory input to excitatory neurons by 19.0±3.3% (mean±s.e.m.), excitatory input to inhibitory neurons by 27.4±3.6%, inhibitory input to excitatory neurons by 60.3±5.7% and inhibitory input to inhibitory neurons by 53.7±5.9% (Figure 1C). ECM depletion did not affect neuronal survival, as indicated by fragmented nuclei quantifications (Figure S3). Because GABAergic synapses were affected more strongly than glutamatergic ones, we concluded that ECM depletion preferentially reduced inhibitory connectivity *in vitro*.

### ECM depletion reduces inhibitory synapse density *in vivo*

Based on these observations, we next analyzed the density of glutamatergic and GABAergic synapses in somatosensory cortex layers 3-5 following ECM depletion *in vivo*. Intracortical injection of ChABC (500 mU in 3 μl 0.1 M phosphate-buffered saline [PBS], 16 hours) unilaterally depleted ECM in the brain, indicated by the absence of PNNs, which in the brain are found around the fast-spiking interneurons expressing a specific potassium channel Kv3.1. The procedure did not affect the density of Kv3.1^+^ neurons and did not alter PNN expression in the contralateral hemisphere (Figure 2A, C). ECM depletion did not influence the density of glutamatergic synapses, but significantly reduced GABAergic synapses in the cortex by 42±6% (Figure 2B, D). Hence, ECM depletion reduced inhibitory connectivity *in vivo*.

**Figure 2.**
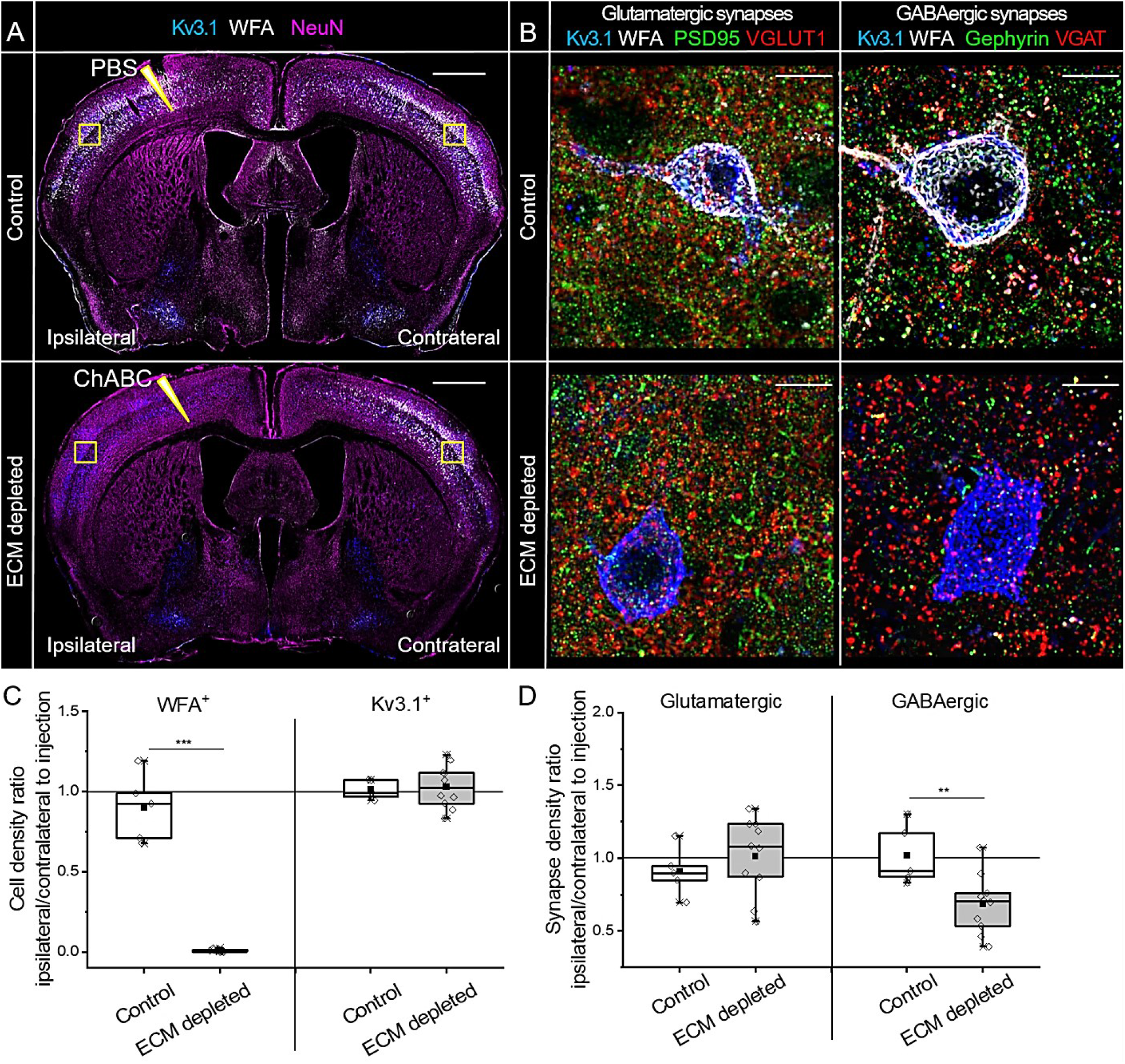
Inhibitory synapse density decreases after ECM depletion *in vivo*. (A) Neuronal nuclei (NeuN, magenta), fast spiking interneurons (Kv3.1, blue) and PNNs (WFA, *Wisteria floribunda* agglutinin, white) were immunohistochemically labeled in brain sections obtained from mice treated with chondroitinase ABC (ChABC, ECM depleted) or phosphate buffered saline (PBS, control) for 16 hours. Sharp triangles indicate intracortical injection sites. Squares indicate the regions in which cell and synapse densities were analyzed. Scale bar, 1 mm. (B) The density of glutamatergic (PSD95-VGLUT1) and GABAergic (Gephyrin-VGAT) synapses was measured in somatosensory cortex layers 3-5. Maximum projections of 56.7×56.7×5 μm regions ipsilateral to the injection sites are shown. Scale bars, 10 µm. (C) Changes in PNN+ and Kv3.1+ neuron densities were quantified as ipsilateral to contralateral ratios. (D) Changes in glutamatergic and GABAergic synapse densities were calculated as ipsilateral to contralateral ratios. Data are shown for each animal examined (n≥5 animals per condition). Data are medians (lines inside boxes)/ means (filled squares inside boxes) ± interquartile ranges (IQR; boxes) with 10/ 90% ranks as whiskers. Open diamonds are data points. The asterisks indicate significant differences with control, based on Kruskal-Wallis tests (***p<0.001, **p<0.01).

### ECM depletion increases the strength of inhibitory synapses

Since ECM depletion reduced the density of inhibitory synapses, we further asked whether the strength of inhibitory input is functionally reduced at the single neuron level. To answer this question, we analyzed spontaneous neurotransmitter release in inhibitory synapses. We measured miniature inhibitory postsynaptic currents (mIPSCs) in mature cultivated neurons using somatic patch clamp recordings (Figure 3A, B).

**Figure 3.**
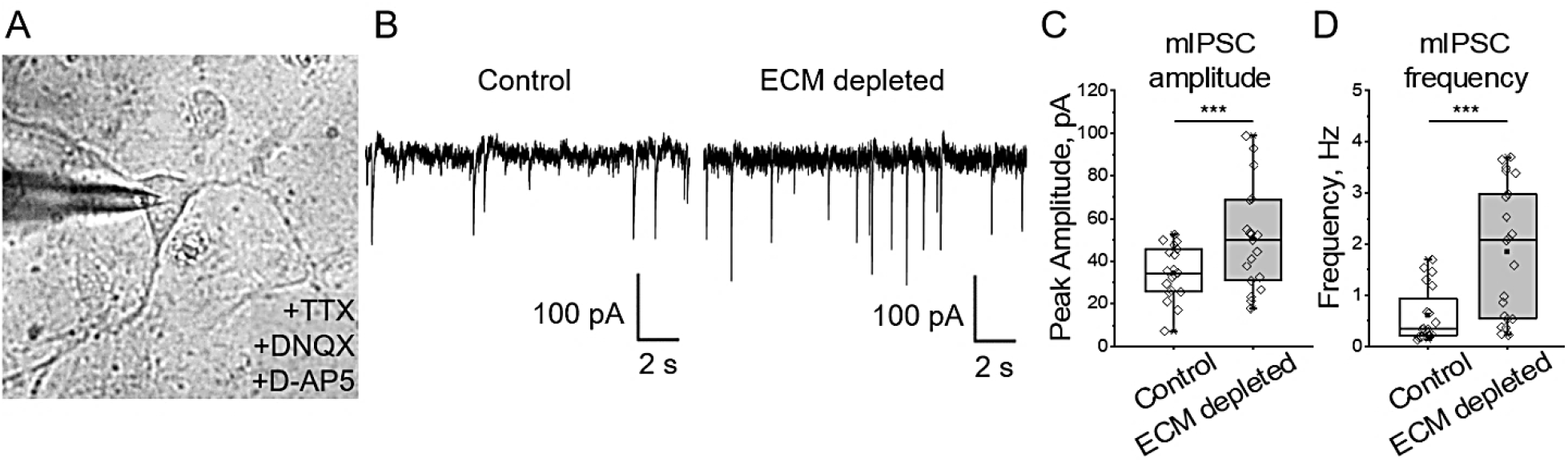
ECM depletion strengthens inhibitory input to single neurons. (A) Somatic patch clamp of a neuron in presence of sodium channel blocker (TTX) and glutamate receptor antagonists (DNQX and D-AP5) reveals miniature inhibitory postsynaptic currents (mIPSCs). (B) Representative current tracks exemplify mIPSCs detected in control and ECM depleted cultures. (C, D) Quantifications of mIPSC amplitude and frequency indicate that ECM depletion increased the total inhibitory input to single neurons (n≥19 neurons per condition, results obtained from 5 independent experiments). Data are medians (lines inside boxes)/ means (filled squares inside boxes) ± IQR (boxes) with 10/ 90% ranks as whiskers. Open diamonds are data points. The asterisks indicate significant differences with control, based on Kruskal-Wallis tests (***p<0.001).

For recording mIPSCs, 1 µM tetrodotoxin (TTX) was applied to prevent action potential-driven synaptic release, and a mixture of glutamate receptor antagonists (10 µM DNQX and 10 µM D-APV) was added to isolate inhibitory currents. Interestingly, ECM depletion increased both amplitude (Figure 3C) and frequency (Figure 3D) of mIPSCs. While the higher mIPSC amplitude indicates elevated neurotransmitter content per synaptic vesicle (Frerking et al., 1995) or increased number and conductance of postsynaptic receptors (Nusser et al., 1997), the higher frequency can result from increased number of synapses and higher release probability (Roberto et al., 2004). Together with the reduction of synapse number (Figure 1C), these data show that ECM depletion increased the strength of inhibitory synapses.

### ECM depletion reduces presynaptic expression of GABA_B_ receptors in inhibitory synapses

To understand how ECM depletion facilitates inhibition, we examined the ultrastructural organization of inhibitory and excitatory postsynapses and analyzed the distribution of GABA receptors. We used stimulated emission depletion (STED) microscopy to uncover the morphology of gephyrin and PSD95 containing scaffolds within structurally complete GABAergic and glutamatergic synapses (Figure 4A).

**Figure 4.**
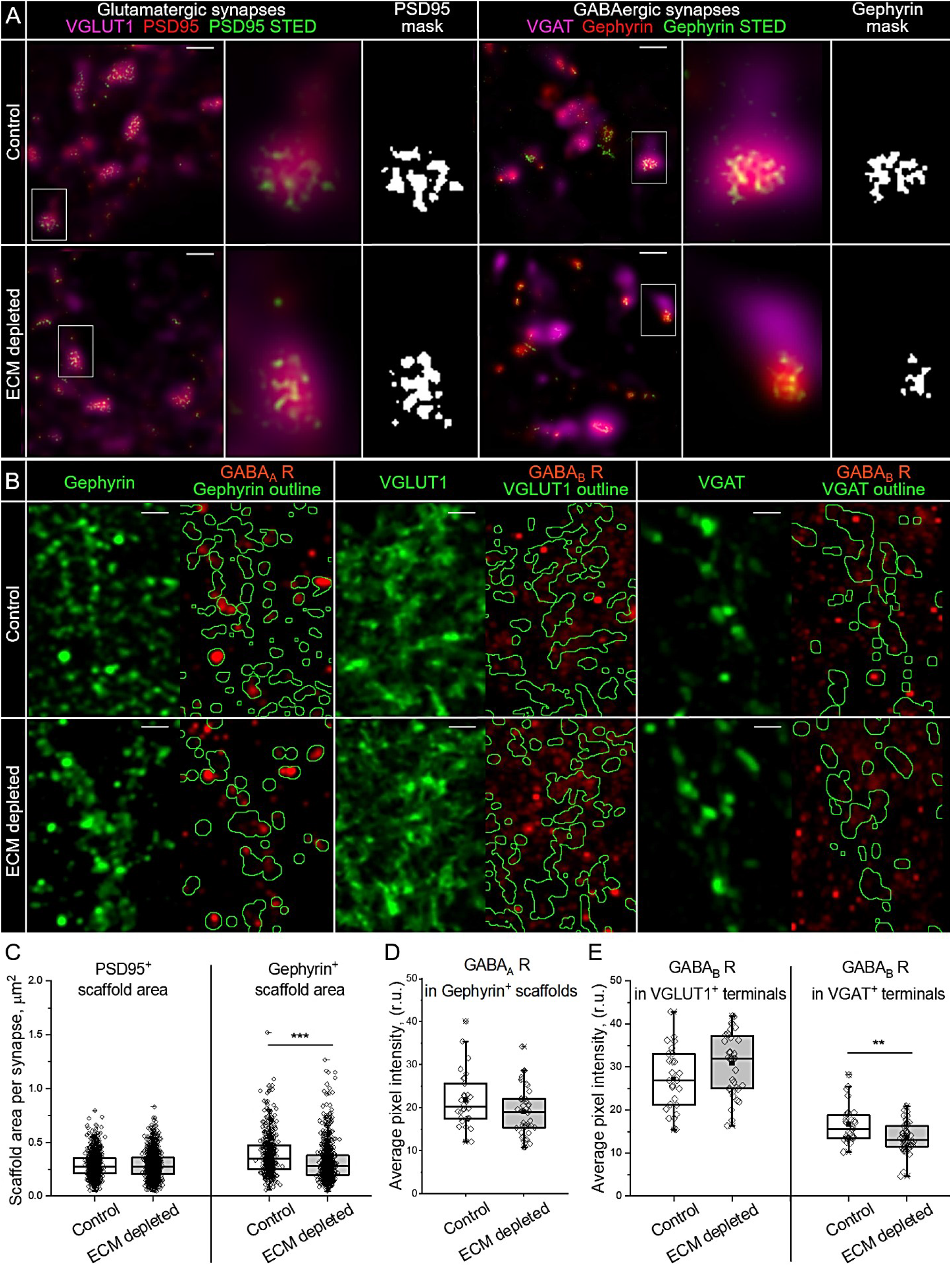
ECM depletion alters the pre- and postsynaptic organization of inhibitory synapses. (A) Stimulated emission depletion (STED) microscopy resolves the morphology of presynaptic scaffolds in glutamatergic and GABAergic synapses. Scale bars, 1 μm. Single synapses highlighted with white rectangles are magnified and the corresponding masks of postsynaptic scaffolds are shown. (B) The panel illustrates the analysis of GABA_A_ receptor (GABA_A_ R) expression in inhibitory postsynapses (gephyrin^+^ areas), GABA_B_ receptor (GABA_B_ R) expression in excitatory (VGLUT1^+^ areas) and inhibitory (VGAT^+^ areas) postsynapses. The outlined areas (green) depict the regions in which the immunoreactivity of GABA receptors was measured. Scale bars, 2 μm. (C) The area of scaffolds containing PSD95 or gephyrin was quantified in single synapses (n≥580 synapses per condition, results from 5 independent experiments). (D) Immunoreactivity of GABA_A_ receptors in GABAergic postsynapses. (E) Immunoreactivity of GABA_B_ receptors in glutamatergic and GABAergic presynapses. (D, E) The average pixel intensity was quantified for each neuron examined (n≥30 cells per condition, results from 5 independent experiments). Data are medians (lines inside boxes)/ means (filled squares inside boxes) ± IQR (boxes) with 10/ 90% ranks as whiskers. Open diamonds are data points. The asterisks indicate significant differences with control, based on Kruskal-Wallis tests (***p<0.001, **p<0.01).

By analyzing the area of binary masks representing single synaptic scaffolds, we determined that ECM depletion reduced the size of gephyrin, but not PSD95 scaffolds (Figure 4C). Because gephyrin scaffolds are essential for the clusterization of postsynaptic GABA_A_ receptors, we measured the immunoreactivity of GABA_A_ receptors inside gephyrin containing postsynapses (Figure 4B, D), but found no significant changes after ECM depletion. The total expression of GABA_A_ receptors was not affected (Figure S3).

GABA_B_ receptors act as negative regulators of neurotransmitter release on the presynaptic side of both excitatory and inhibitory synapses. By measuring the immunoreactivity of GABA_B_ receptors on VGLUT1^+^ and VGAT^+^ presynaptic terminals, we revealed that ECM depletion preferentially reduces GABA_B_ receptor expression in GABAergic presynapses (Figure 4 B, E). These results suggest that the increased strength of inhibitory synapses after ECM depletion is associated with reduced reciprocal inhibition of neurotransmitter release. Neither total nor postsynaptic expression of GABA_B_ receptors was significantly influenced by ECM depletion (Figures S4 and S5).

### ECM depletion increases neuronal activity and facilitates spiking-bursting transitions

Knowing that ECM depletion reduces inhibitory connectivity, but enhances inhibitory input to single neurons, we asked how these opposing changes affect neuronal activity at the network level. Spontaneous network activity was investigated using multiple electrode arrays (MEAs). Within this methodology, neuronal cultures are grown on an array of electrodes, each detecting population spikes and bursts, generated by small groups of neurons (10-15 in our experiments), located within 100 µm radius from the center of the electrode (Figure 5A, B). While the frequency of spikes measured as the electrode mean firing rate (MFR) represents general neuronal activity, the network phase transitions are represented by bursting behavior, which was evaluated as mean bursting rate (MBR) changes. In mature networks, spiking-bursting transitions are mostly synchronized, as indicated by the alignment of burst events (Figure 5C). Therefore, MBR measurements also partially reflect neuronal network synchrony.

**Figure 5.**
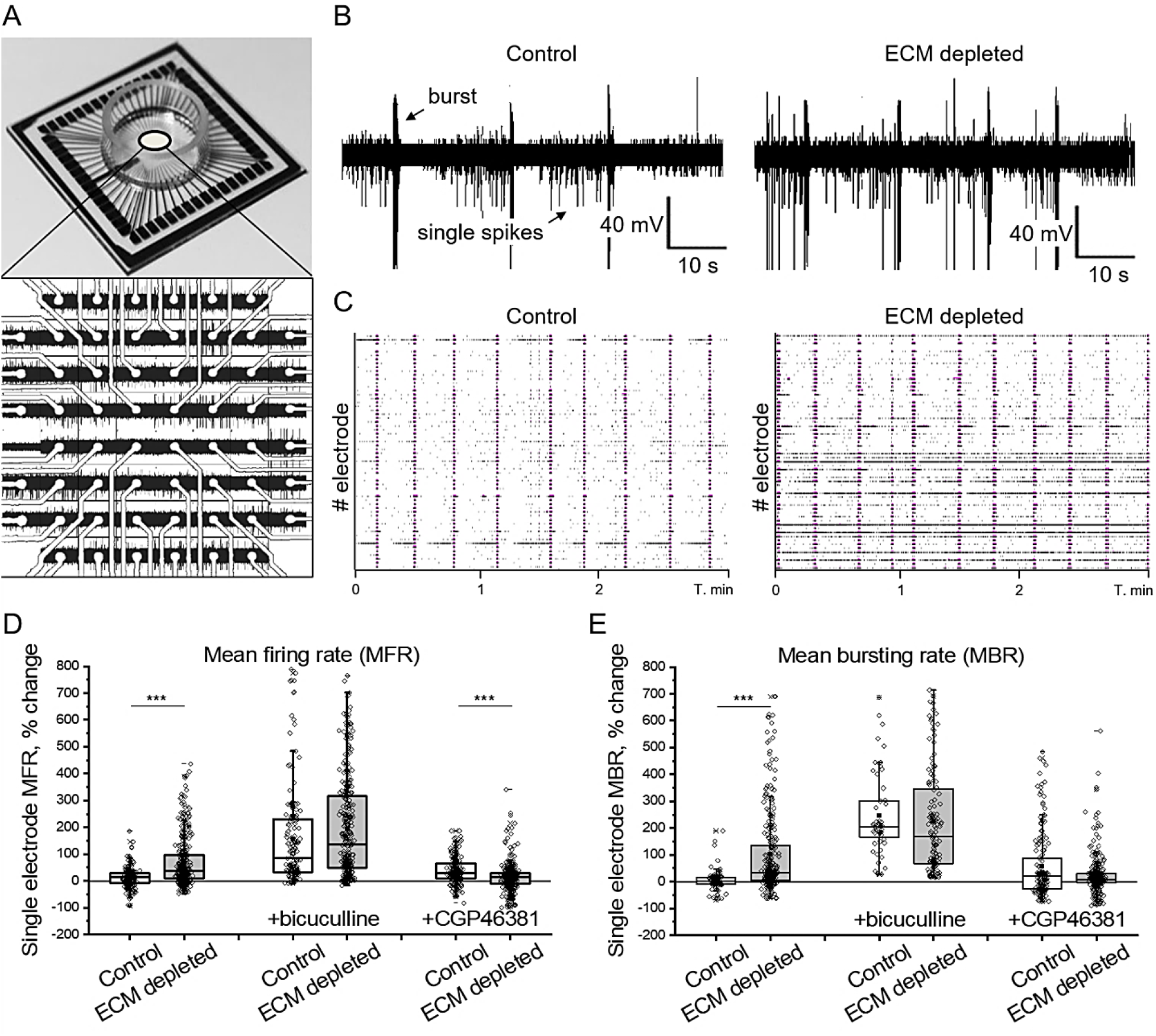
The increase of neuronal network activity after ECM depletion is inhibition-dependent. (A) Neuronal network activity was examined using multiple electrode arrays (MEAs). The panel demonstrates the layout and network activity recorded on a MEA chip with a square array of 59 electrodes. (B) Representative voltage tracks exemplify spikes and bursts detected by single electrodes in control and ECM depleted cultures. (C) Raster plots show synchronized network activity in control and ECM depleted cultures. Black ticks are single spikes, magenta bars are burst events. The changes of (D) mean firing rate (MFR) and (E) mean bursting rate (MBR) were quantified for single electrodes as the differences with the baseline activity of the same electrode before treatment (n≥169 electrodes per condition, results from 5 independent experiments). The effects of GABA_A_ and GABA_B_ receptor blockage were analyzed by comparing single electrode activity before and after incubation with the antagonist (6 μM bicuculline and 100 μM CGP46381, respectively). Data are medians (lines inside boxes)/ means (filled squares inside boxes) ± IQR (boxes) with 10/ 90% ranks as whiskers. Open diamonds are data points. The asterisks indicate significant differences with control, based on Kruskal-Wallis tests (***p<0.001).

We analyzed how ECM depletion alters neuronal activity and bursting behavior by measuring MFR and MBR changes after ChABC (500 mU/ml, 16 hours) or control (PBS, 16 hours) treatment. The activity recorded by single electrodes was compared to the baseline activity of the same electrode before treatment (Figure 5D, E). On average, ECM depletion increased MFR by 69.8±5.2% (mean±s.e.m.) and MBR by 102.0±9.6% (mean±s.e.m.). The increase of neuronal network activity and facilitated bursting was inhibition-dependent, since the blockage of GABA_A_ receptors (6 µM bicuculline metiodide for 30 minutes) levelled the effect of ECM depletion. The blockage of GABA_B_ receptors (100 µM CGP46381 for 30 minutes) moderately enhanced neuronal activity. After ECM depletion, the effect of GABA_B_ receptor blockage was significantly reduced, as indicated by MFR quantification after treatment with CGP46381. Hence, the increase of neuronal network activity after ECM depletion functionally involved the downregulation of GABA_B_ receptors.

### Reduction of inhibitory connectivity after ECM depletion outweighs inhibitory synapse strength increase

The number and strength of GABAergic synapses are essential determinants of inhibitory input that control the activity and synchronization in neuronal networks. We revealed that these parameters undergo opposing changes after ECM depletion, implying the necessity to understand their interaction at the network level. Using the earlier elaborated method (Dzyubenko et al., 2017), we reconstructed the observed alterations *in silico* (Figure 6A).

**Figure 6.**
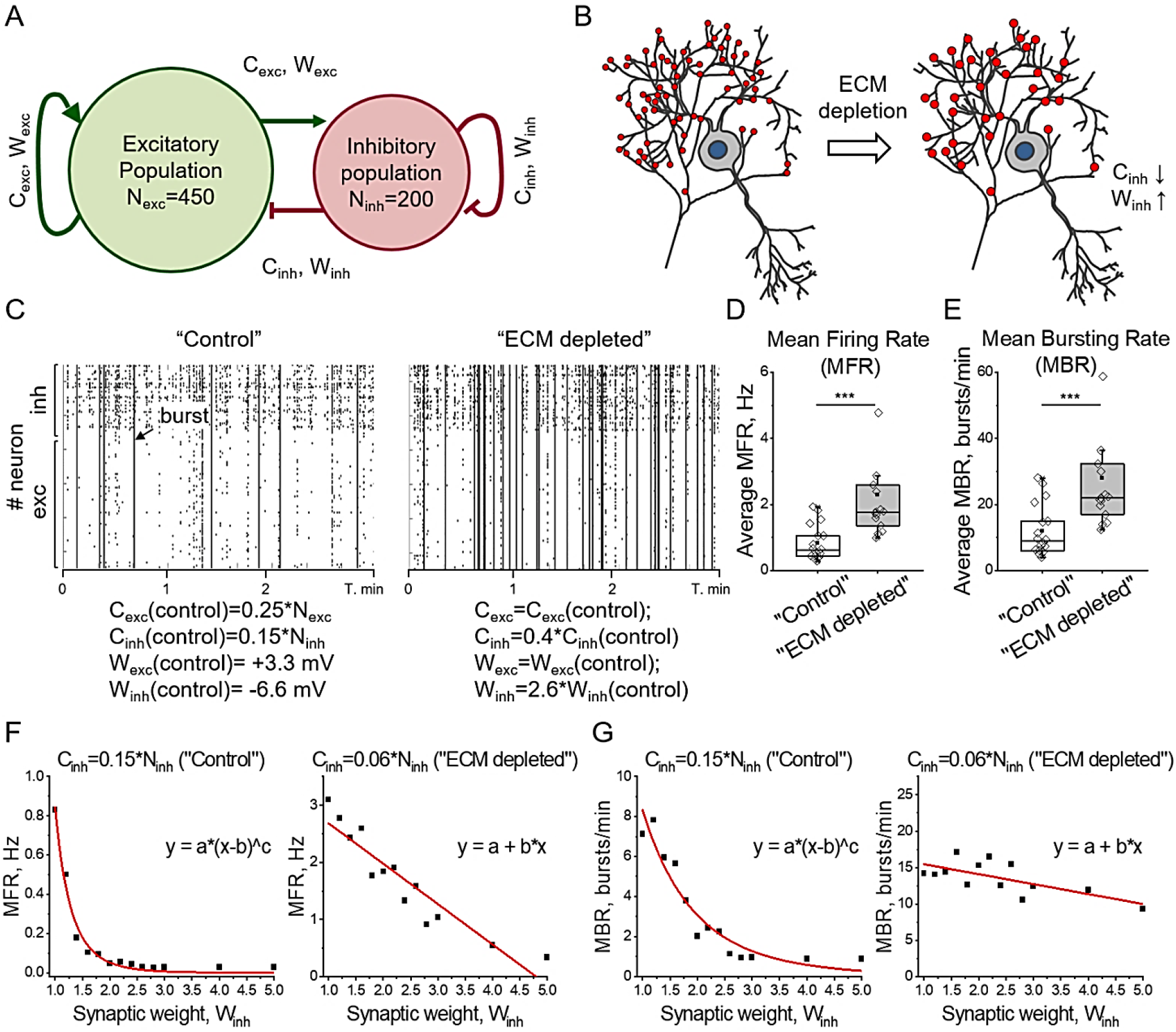
Network activity simulation *in silico* indicates the prevailing role of inhibitory connectivity reduction following ECM depletion. (A) The schematic drawing illustrates the model of the spiking neuron network. Ni_nh_ and N_exc_ are numbers of inhibitory and excitatory neurons, Winh and W_exc_ are weights of corresponding synapses. (B) The ECM depletion was mimicked by tuning C_inh_ and Winh parameters in accordance with experimentally observed changes. (C) Raster plots exemplify the activity of “control” and “ECM depleted” networks. The corresponding simulation parameters are depicted. Ticks are single spikes, vertical dashes indicate burst events. The quantification of network average (D) mean firing rate (MFR) and (E) mean bursting rate (MBR) is shown for “control” and “ECM depleted” simulation conditions. Data are medians (lines inside boxes)/ means (filled squares inside boxes) ± IQR (boxes) with 10/ 90% ranks as whiskers. Open diamonds are data points. The asterisks indicate significant differences with the control, based on Kruskal-Wallis tests (***p<0.001). (F) MFR and (G) MBR are quantified in a range of Winh changes for “control” and “ECM depleted” inhibitory connectivity. Note that the reduction of inhibitory connectivity switches the dependence of network activity on inhibitory synapse weight from power law to linearity. Squares indicate the mean of simulation repetitions, fit functions are shown in red. For each condition, 15 independent simulation experiments were performed.

Excitatory and inhibitory connectivity was defined as the proportion of all neurons providing the input to a single cell (C_exc_=0.25*N_exc_, C_inh_=0.15*N_inh_ in “control”), based on the previous study (Izhikevich, 2003) and connectivity estimations in neuronal cultures (Pastore et al., 2018). The strength of single connections was defined by synaptic weights reflecting membrane potential changes upon synapse activation (W_exc_=+3.3 mV, W_inh_=-6.6 mV in “control”). Thereby, excitation and inhibition were balanced, and the computations were performed in a near-critical state (Figure S6) characterized by stable spiking-bursting transitions. ECM depletion was simulated by modifying inhibitory connectivity (C_inh_) and synaptic weights (W_inh_) in accordance with experimentally observed changes (Figure 6B). In agreement with *in vitro* experiments, ECM depletion increased network MFR (Figure 6C, D) and bursting rates (Figure 6C, E). Of note, our approach closely resembled the intrinsic variability in real neuronal networks, because the connectivity matrix was newly generated for each simulation instance.

To compare the impact of C_inh_ and W_inh_ on the resulting neuronal activity, we measured MFR and MBR over a range of different C_inh_ and W_inh_ values (Figure S7). Excitatory input parameters C_exc_ and W_exc_ were set to control values. The reduction of C_inh_ negatively and linearly correlated with network MFR (r=-0.98; p<0.01) and MBR (r=-0.92; p<0.01), while the increased C_inh_ strongly diminished neuronal activity. Under moderate decrease of C_inh_ (C_inh_=0.125*N_inh_ and C_inh_=0.1*N_inh_), increasing W_inh_ partially compensated MFR and MBR changes. However, when C_inh_ was set in accordance with ECM depletion effect *in vitro* (40% of control, C_inh_=0.06*N_inh_), both MFR and MBR remained elevated in a broad range of W_inh_. While changing inhibitory connectivity from C_inh_=0.15*N_inh_ in “control” simulation to C_inh_=0.06*N_inh_ in “ECM depleted” simulation, the dependence of MFR and MBR on W_inh_ switched from a power law to linearity (Figure 6F, G). Conclusively, the reduction of inhibitory connectivity after ECM depletion outweighed the strengthening of inhibitory synapses and increased the resulting activity of neuronal networks.

## Discussion

Here we demonstrate that the brain ECM supports the maintenance of neuronal network E-I balance by retaining inhibitory connectivity. ECM depletion preferentially decreases the density of inhibitory synapses and the size of inhibitory postsynaptic scaffolds, while it homeostatically increases inhibitory synapse strength. Commonly, inhibitory synapse scaling downregulates neuronal network activity and synchronization, providing a key mechanism for neuronal activity adjustment (Sprekeler, 2017). After ECM depletion, the degree of inhibitory connectivity reduces to an extent that inhibitory synapse scaling is no longer efficient in controlling the state of neuronal networks. As a result, neuronal network activity and synchrony increase.

We observed that ~60% of GABAergic synapses are lost already 16 hours after ECM depletion. How could such a major decrease in synapse density occur without increased neuronal cell death? Recent findings indicate that neuronal networks are continuously remodelled and that synapses dynamically wane and re-emerge within a few days (Pfeiffer et al., 2018). Inhibitory synapses are especially dynamic, about 60% of them retract and return at the same place within 4 days (Villa et al., 2016). Intuitively, this structural volatility but spatial persistency implies the existence of a stable framework to secure the integrity of neuronal circuits. Our data suggests that brain ECM provides such framework to stabilize inhibitory connectivity. After ECM depletion, the reduction of GABAergic synapse density may be due to the increased volatility and inability of synaptic boutons to find their postsynaptic site. The demarcation of postsynaptic sites has been proposed to be defined by PNNs (Fawcett et al., 2019; Sigal et al., 2019), the facet-like ECM coatings that compartmentalize neuronal surface. On neurons devoid of PNNs, a less dense ECM layer may potentially play a similar role. An increased ratio of excitatory to inhibitory synapses has recently been shown in the hippocampus of mice exhibiting simultaneous knockout of several ECM proteins and glycoproteins as a consequence of disturbed PNN formation (Gottschling et al., 2019).

At the level of single synapses, ECM depletion increases inhibitory synapse strength, as indicated by the increased mIPSC amplitude and frequency together with the decreased synapse density. With STED microscopy, we observed the reduced size of postsynaptic gephyrin scaffolds after ECM depletion. However, despite the known role of gephyrin for postsynaptic GABA_A_ receptor clustering and stabilization (Choii and Ko, 2015), neither expression nor localization of GABA_A_ receptors was affected. Apparently, the gephyrin containing scaffolds condensed and retained GABA_A_ receptors. On the presynaptic side, we found a significant reduction of GABA_B_ receptor expression on VGAT^+^ terminals, which functionally associated with the diminished sensitivity to the GABA_B_ antagonist CGP46381. Presynaptic GABA_B_ receptors act as activity-dependent regulators of neurotransmitter release. Upon repetitive stimulation, the spillover of GABA activates presynaptic GABA_B_ receptors, transiently reducing subsequent neurotransmitter release (Davies et al., 1990; Pitler and Alger, 1994). Our results indicate that ECM depletion attenuated this mechanism and increased inhibitory synapse strength via presynaptic facilitation. It remains unclear how exactly ECM regulates GABA_B_ receptor localization and function. The extracellular complement control protein module CCP1 of GABA_B_ R1 subunit interacts with laminin α5 subunit and fibulin-2 of ECM (Blein et al., 2004; Pless, 2009), but the functional consequences of this interaction need further investigation.

At the network level, the reduction of inhibitory connectivity outweighs the increased inhibitory synapse strength after ECM depletion. Thereby, the E-I balance switches towards excitation, resulting in the increased neuronal activity and network synchrony, as evidenced by MEA recordings and *in silico* simulations. A similar increase of network activity could arise from the reduced excitatory input to GABAergic interneurons, as observed in the visual cortex after ECM depletion (Faini et al., 2018), and the decreased rate of their spontaneous firing, as observed in a model of peritumoral epilepsy (Tewari et al., 2018). Unlike the VGLUT2^+^ thalamic inputs in the visual cortex (Faini et al., 2018; Nahmani and Erisir, 2005), ECM depletion does not significantly alter the local excitatory connectivity in somatosensory cortex layers 3-5, as indicated by the VGLUT1^+^ synapse quantifications. We therefore conclude that controlling the density of excitatory synapses on GABAergic interneurons is not likely to be a key mechanism by which extracellular matrix supports excitation-inhibition balance in local neuronal networks. However, the decreased rate of spontaneous firing of GABAergic interneurons may indeed amplify the E-I balance changes by further decreasing the efficiency of inhibitory control over neuronal network activity.

Altered E-I balance following ECM breakdown is likely a key component of pathophysiology of psychosis (Pantazopoulos et al., 2015; Soleman et al., 2013) and epilepsy (Arranz et al., 2014; Tewari et al., 2018). In ischemic stroke, adjusting the E-I balance after ECM decomposition may, on the contrary, support neurological recovery. Post-stroke neuroplasticity is impaired by decreased neuronal excitability in perilesional brain areas (Clarkson et al., 2010; Wang et al., 2018). In light of the new evidence we present here, the transient decline in cortical ECM integrity after ischemia (Dzyubenko et al., 2018) may support neuronal network rewiring by stimulating neuronal activity. Hence, the controlled degradation of ECM could be a promising target in stroke therapy, as it may allow promoting neuronal activity and plasticity.

The therapeutic potential of ECM degradation in the injured brain will crucially depend on the precise targeting of ECM modifications. The crude ablation of ECM elicits memory loss and learning deficits (Hylin et al., 2013; Shi et al., 2019). In this study, we demonstrate that the near-complete ECM digestion disrupts criticality in neuronal networks, indicated by the switch of dependence between neuronal activity and inhibitory synapse strength from a power law to linearity. The dynamic tuning of cortical circuits to criticality is essential for efficient information processing in the brain (Gautam et al., 2015; Kinouchi and Copelli, 2006; Ma et al., 2019). Refined tools for controlled ECM decomposition will not only expand opportunities for research but will also open new directions in neurorestorative therapies.

## Materials and methods

### Legal issues and animal housing

Experimental procedures were approved by the local government (Bezirksregierung Düsseldorf) and conducted in accordance to European Union (Directive 2010/63/EU) guidelines for the care and use of laboratory animals. C57BL/6j mice (Envigo, Indianapolis, IN, U.S.A.) were kept in groups of 5 animals/cage in a regular inverse 12 h light-dark cycle and access to food and water *ad libitum*. All efforts were made to reduce the number of animals in the experiments.

### Cell cultures

Primary cultures of neurons and astrocytes were prepared as described previously (Gottschling et al., 2016). Hippocampal neurons were obtained at embryonic day 15 (E15) and cortical astrocytes were obtained at postnatal day 1 (P1) from male and female mice. Neurons were supported by astrocyte monolayers cultivated on cell culture inserts with a permeable membrane (Figure S1A), allowing for the long term maturation of neuronal networks. We plated 50,000 neurons in a 50 µl droplet onto pre-treated glass coverslips or MEA chips. Cell culture inserts containing 50,000 astrocytes were combined with neuronal cultures on the same day. The cultures were maintained in Neurobasal medium (21103049, ThermoFisher, Waltham, MA, U.S.A.) supplemented with 2 mM L-glutamine (25030081, ThermoFisher), 1% v/v B27 (A3582801, ThermoFisher) and 1% v/v SM1 (05711, STEMCELL Technologies, Vancouver, Canada). We changed half of the medium weekly to keep the pH around 6.5. All *in vitro* experiments were performed on fully mature neurons after 21-28 days of cultivation. The evolution of neuronal networks *in vitro* was characterized by coherent maturation of synaptic connectivity and ECM expression, partially resembling neuronal circuit development *in vivo* (Choi, 2018). In the course of cultivation, synapse density increased until maturity was reached after 21 days in vitro (DIV) (Figure S1B, C). The establishment of network connectivity correlated with the expression of PNNs, the condensed ECM layers characteristic for the mature neuronal cultures (Figure S1B, D). The network activity in neuronal cultures evolved accordingly, changing from the high frequency random spiking at 7 DIV followed by the low activity period at 14 DIV and the regular bursting pattern at 21 DIV (Figure S1E). At later time points (i.e., 35 DIV) synapse density, PNN expression and network activity were stabilized and did not change. Similar to our previous work (Dzyubenko et al., 2017), the mature neuronal cultures contained 34±6.3% (mean±s.e.m.) inhibitory interneurons (Figure S2).

### ECM depletion

We followed an established approach for enzymatic ECM digestion (Bikbaev et al., 2015; Pizzorusso et al., 2002). Cell cultures were incubated with 500 mU/ml ChABC (C3667, Sigma-Aldrich, Taufkirchen, Germany) or 500 U/ml hyaluronidase (HYase, H4272, Sigma-Aldrich) for 16 hours. For ECM depletion *in vivo*, 500 mU ChABC or 500 U HYase were dissolved in 2 µl of 0.1 M PBS and delivered via a single stereotactic (bregma 0, left 3 mm, deep 1 mm) intracortical injection. At 16 hours post injection, animals were sacrificed and brains were processed for further analysis. Control animals were treated with vehicle (0.1 M PBS). In our hands, the two enzymes were equally efficient for ECM digestion and specifically targeted the neuronal cell culture compartment, while no alterations in glial cells were detected (Figure S7). Moreover, ChABC and HYase induced identical changes in synapse density and network activity (Figure S9).

### Immunolabelling procedures

For immunohistochemistry, brains were perfused with 4% w/v paraformaldehyde (PFA) and post-fixed for 12 hours in 4% w/v PFA. 40 µm coronal free-floating sections were obtained from the bregma level. For immunocytochemistry, cell cultures were fixed with 4% w/v PFA for 10 min at room temperature. Synaptic proteins were detected mouse anti-PSD95 (1:500, MAB1598, Millipore, Burlington, MA, U.S.A.), guinea pig anti-VGLUT1 (1:500, 135304, Synaptic Systems, Goettingen, Germany), mouse anti-gephyrin (1:500, 147011, Synaptic Systems) and guinea pig anti-VGAT (1:500, 131103, Synaptic Systems) antibodies. For GABA receptor quantification, GABA_A_ and GABA_B_ receptors were labelled with chicken anti-GABA_A_ γ^2^ (1:500, 224006, Synaptic Systems) and rabbit anti-GABA_B_ (1:500, 322102, Synaptic Systems) antibodies. To characterize ECM expression, we applied biotinylated WFA (1:100, B-1355, Vector Laboratories, Burlingame, USA), biotinylated hyaluronan binding protein (1:100, 400763, AMS Biotechnology, Frankfurt, Germany), rabbit anti-aggrecan antibody (1:500, AB1031, Millipore) and rat anti-473HD antibody (1:100, produced by the group of Prof. Andreas Faissner, Bochum, Germany; (von Holst et al., 2006)). Neuronal types were identified using rabbit anti-GABA (1:2000, A2052, Sigma-Aldrich), chicken anti-NeuN (1:300, ABN91, Millipore), mouse anti-neurofilament M (1:500, 171231, Synaptic Systems) and rabbit anti-Kv3.1b (1:1000, APC-014, Alomone Labs, Jerusalem, Israel) antibodies. Rat anti-glial acidic fibrillary protein (GFAP; 1:1000, 13-0300, ThermoFisher), mouse anti-β-catenin (1:500, ab19381, Abcam, Cambridge, UK) and rabbit anti-connexin 43 (1:500, 3512, Cell Signaling Technologies, Frankfurt, Germany) antibodies were used as astroglial markers. For fluorescence detection, secondary antibodies conjugated to Alexa or Atto dyes were used. Nuclei were counterlabeled with DAPI (1:1000, D1306, ThermoFisher).

### Low-resolution microscopy for basic quantifications

For basic quantifications of cell density and marker proteins expression following immunohistochemistry/ immunocytochemistry, four 425.1×425.1 μm regions of interest (ROIs) per condition per experiment were selected at random positions for cell culture specimens. In brain sections, the ROIs were positioned in the left and right cerebral somatosensory cortex layers 3-5. Single plane micrographs were obtained using the Carl Zeiss LSM710 confocal microscope (20x Plan Apochromat objective, NA 0.8, pixel size 0.21 μm).

### Total expression of GABA receptors

The total expression of GABA receptors on the neuronal surface was investigated *in vitro* by Western blot. Membrane proteins were extracted using the Mem-PER Plus Membrane Protein Extraction Kit (89842, ThermoFisher) and separated by sodium dodecyl sulfate polyacrylamide gel electrophoresis (SDS-PAGE) on 1 mm 8% polyacrylamide gels. To avoid protein aggregation the samples were not heated. The proteins were transferred onto nitrocellulose membranes using the Trans-Blot Turbo Transfer System (Biorad, Hercules, CA, U.S.A.) mixed molecular weight program, followed by pre-blocking in 3% bovine serum albumin (BSA) for 1 hour. Then, the membranes were incubated with primary chicken anti-GABA_A_ γ^2^ (1:1000, 224006, Synaptic Systems), rabbit anti-GABA_B_ (1:1000, 322102, Synaptic Systems) and mouse anti-synaptotagmin-1 (1:1000, 105011, Synaptic Systems) antibodies for 72 h at 4°C. Secondary antibodies were applied stepwise, and the proteins were visualized in separated fluorescence and luminescence channels. For fluorescence detection, Alexa-647 and Cy-3 conjugated antibodies were used. Chemiluminescence was detected with HRP conjugated antibodies using Pierce ECL Western Blotting-Substrate (32106, ThermoFisher). Multiple proteins were detected on the same membrane using the ChemiDoc XRS+ Imaging System (Biorad), and labelling intensity was normalized to the stain-free signal. The molecular weights of the proteins were verified using a prestained protein ladder (ab116028, Abcam). Data were analyzed by densitometry in ImageJ using the gel quantification plugin.

### Synapse density and synaptic GABA receptor quantifications

The density of glutamatergic and GABAergic synapses was quantified using a previously established method (Dzyubenko et al., 2016b). For synapse analysis *in vitro*, five 66.5×66.5×5 µm ROIs per condition per experiment were selected at random positions, containing the soma and proximal dendrites of single neurons. In brain sections, five 51×51×10 µm ROIs per condition per experiment were selected in the left and right cerebral cortex layers 3-5. The confocal stacks were obtained using the LSM710 confocal microscope (100x alpha Plan-Apochromat objective, NA 1.46; Carl Zeiss, Jena, Germany). Structurally complete synapses were identified by the overlapping immunolabelling of pre- and postsynaptic markers, which were analyzed with an in-house Synapse Counter plugin for ImageJ (freely available at https://github.com/SynPuCo/SynapseCounter). Therefore, our synapse quantification approach detects the majority of inputs, which a particular cell receives from its local network partners. The expression of pre- and postsynaptic GABA_B_ and GABA_A_ receptors was evaluated by immunofluorescence intensity analysis. Pre- and postsynaptic structures were analyzed using Synapse Counter, and mean pixel intensities were determined as estimates of protein expression changes.

### STED microscopy of postsynaptic scaffolds

The morphology of postsynaptic scaffolds was investigated by STED microscopy using a previously established method (Dzyubenko et al., 2016b). We employed the time-gated Leica TCS SP8 microscope (Wetzlar, Germany), which is equipped with white light pulse laser (WLL2) and gated hybrid detection. An oil immersion HCX PL APO STED 100x (numerical aperture 1.4) objective was used. The ultrastructure of postsynaptic scaffolds within structurally complete synapses was analyzed. First, 23.8×23.8 µm single-plane confocal images were obtained using 488 and 633 nm excitation wavelengths for the post- and presynaptic markers, respectively. Then, the STED scans were obtained in the same stage position using 488 nm excitation and 592 nm depletion lasers. The detection time-gating interval was set to 6-10 ps post-pulse time window. All settings were kept constant throughout the experiments. To improve the visibility of fine structural elements, the raw data were deconvolved using Hyugen’s software. The binary masks of single synaptic scaffolds were generated using automated thresholding (Otsu method) in ImageJ, and the mask area was measured.

### Whole-cell mIPSC recordings

Neuronal cultures were recorded in patch-clamp whole-cell configuration using Axopatch 200B amplifier (Molecular Devices, San Jose, CA, U.S.A.) with pClamp software 10.6. Microelectrodes of 1.5 mm thin-walled filamented borosilicate glass (World Precision Instruments, Friedberg, Germany) were pulled with a DMZ-Universal Puller (Zeitz Instruments, Martinsried, Germany) and polished to a final resistance of 3–4 MΩ. Neurons were voltage-clamped at −60 mV, the signals were filtered at 1.0 kHz and recorded with 10 kHz. Series resistance and cell capacitance were compensated prior to the recordings. BrainPhys basal medium was used as extracellular solution. The pipette solution contained 140 mM KCl, 1 mM CaCl_2_·2H_2_O, 4 mM MgCl_2_, 10 mM HEPES, 0.4 mM Na_2_-GTP, 4 mM Mg-ATP and 10 mM EGTA (pH 7.3). For the recording of mIPSCs we applied TTX (1 µM) to prevent action potential-driven synaptic release. The inhibitory postsynaptic currents were pharmacologically isolated using glutamate receptor antagonist DNQX (10 µM) and NMDA receptor antagonist D-APV (10 µM). Data were analysed with Clampfit software 10.6 (Molecular Devices).

### Spontaneous network activity recordings

Spontaneous activity in neuronal networks was measured using cell culture compatible square 8×8 electrode MEA (60MEA200/30iR-Ti, Multi Channel Systems, Reutlingen, Germany), on which primary neurons were grown. To evaluate the effects of ECM depletion, we recorded the baseline activity prior to the application of digesting enzymes, and compared it with the post-treatment activity after 16 hours. The control cultures were treated with 0.1 M PBS. To study the impact of ECM depletion on GABAergic neurotransmission, GABA_A_ (6 µM bicuculline metiodide, Tocris, Bristol, UK) or GABA_B_ (100 µM CGP46381, Tocris) receptor antagonists were applied. After 30 minutes of incubation with the antagonist, network activity was recorded. The effects of GABA antagonists were evaluated with reference to neuronal activity after ECM depletion or control treatment. In all experiments, we recorded spontaneous network activity for 15 minutes (temperature stabilized at 35°C, gas exchange prevented) using the MEA2100 60-channel headstage with the sampling frequency of 40,000 Hz using MC Rack. For each electrode, the mean firing rate (MFR) and mean bursting rate (MBR) were analyzed in MatLab with the SpyCode toolbox generously provided by Dr. Michela Chiappalone (Bologna et al., 2010).

### Network activity simulations

To evaluate the impact of connectivity alterations versus the synaptic strength changes induced by ECM depletion, we implemented a computational approach that we previously established (Dzyubenko et al., 2017). Experimentally observed effects of ECM depletion were reconstructed using an *in silico* network of spiking neurons. The key variables of our model define neuronal response properties and network connectivity. The physiology of fast spiking interneurons and primary excitatory neurons was replicated using previously defined parameters (Izhikevich, 2003). Network circuitry was defined by the sparseness of connectivity (C_exc_, C_inh_) and synaptic weights (W_exc_, W_inh_). Sparseness of connectivity was defined as the proportion of all neurons of a certain type providing the input to a single cell on average: C_exc_=0,25*N_exc_ for excitatory, C_inh_=0,15*N_inh_ for inhibitory input in control condition. Synaptic weights were set as absolute values of postsynaptic membrane potential changes after activation of a synapse. We modified C_inh_ and W_inh_ within a range of biologically feasible values to determine their impact on neuronal network activity. For each simulation instance, network connectivity was newly generated, mimicking the intrinsic variability of real neuronal networks.

### Statistics

For non-normally distributed datasets, data were evaluated by Kruskal-Wallis tests using OriginPro2020 software. For multiple comparisons, Bonferroni correction was applied. Data were presented as box plots depicting the medians (lines inside boxes)/ means (filled squares inside boxes) ± IQR (boxes) with 10% and 90% ranks as whiskers. For normally distributed datasets, e.g. the *in silico* simulations, data were evaluated by two-tailed independent Student’s t-tests. Data were shown as mean±s.e.m. columns. Data points were indicated as diamonds. P values <0.05 were defined to indicate statistical significance.

## Supplement

**Figure S1.**
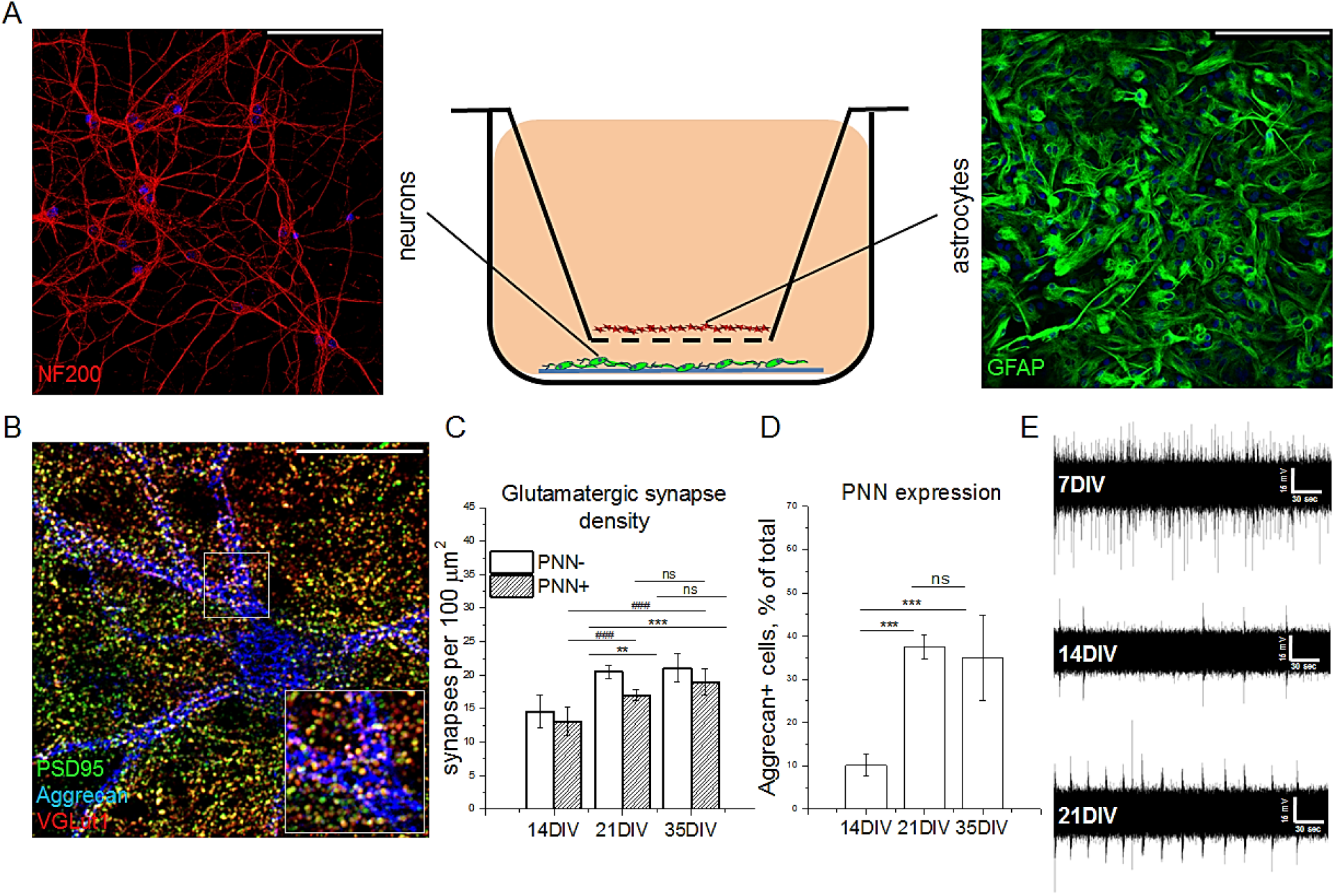
Coherent maturation of synaptic connectivity, extracellular matrix and neuronal network activity in primary neuron-astrocyte co-cultures. (A) The panel illustrates the principle of the indirect co-cultivation of primary neurons and astrocytes. It this setup, the two cell types do not contact each other directly, but share the same medium. Scale bars, 100 µm. (B) The representative micrograph of a single PNN coated neuron is shown. The major proteoglycan of PNNs is labelled with specific anti-aggrecan antibody (blue). Glutamatergic synapses are labeled with anti-PSD95 (green) and anti-VGLUT1 (red) antibodies. The square inlet illustrates that synapse formation predominantly occurs in the areas devoid of aggrecan labelling. Scale bar, 30 µm. The density of structurally complete glutamatergic synapses (C) and the percentage of PNN expressing neurons (D) is quantified after 14, 21 and 35 days of cultivation (DIV). The results of quantification (n≥24 ROIs measuring 66.5×66.5 µm, results obtained from 4 independent experiments) are expressed as mean±s.e.m. Statistically significant differences are indicated with asterisks and hashes, based on Kruskal-Wallis tests (***, ###p<0.001; **,##p<0.01). (E) The evolution of neuronal activity detected by a single electrode is shown over the time of cultivation.

**Figure S2.**
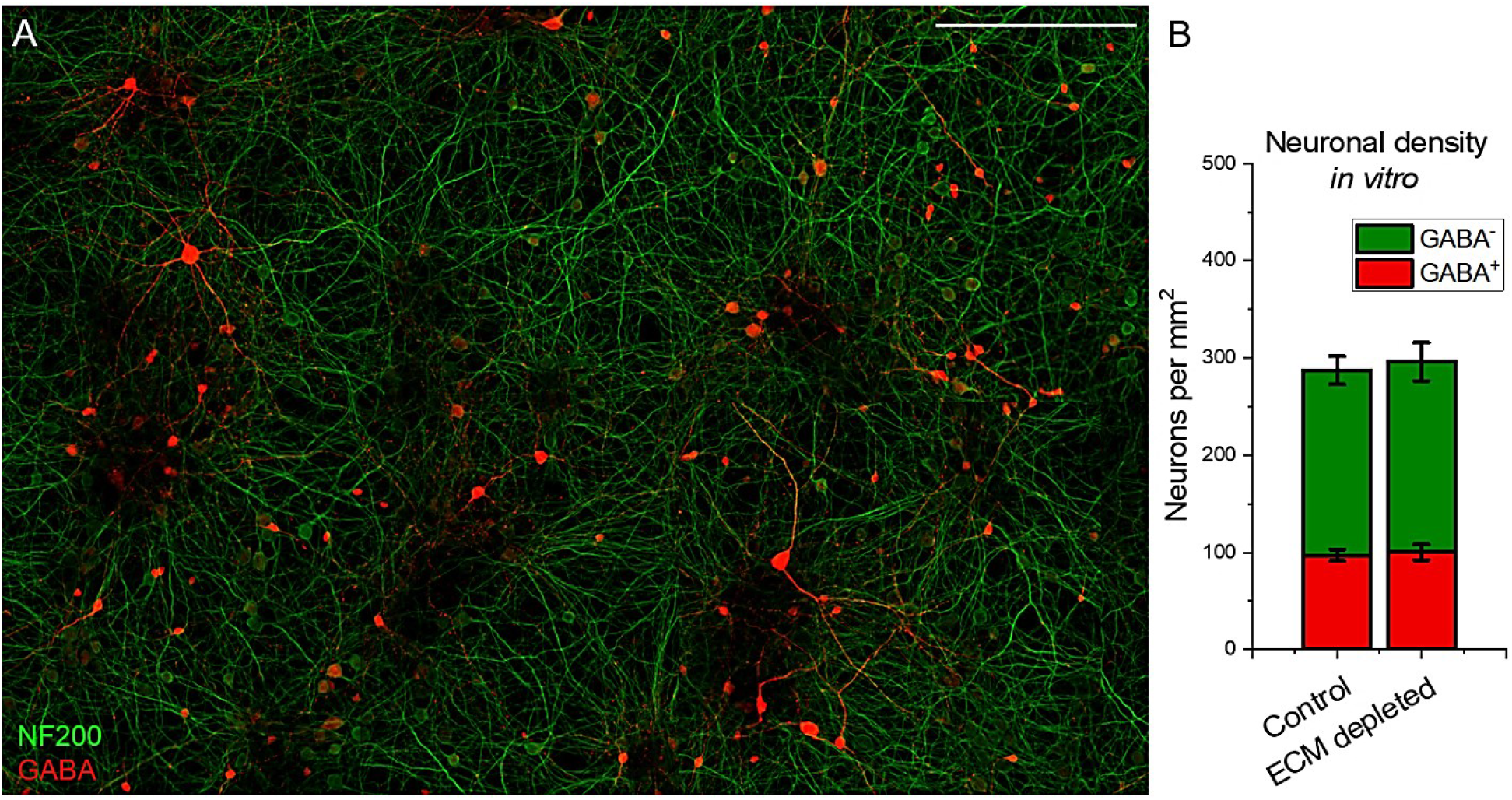
Primary neuronal culture after 21 days of cultivation. (A) The micrograph shows representative labelling of neuronal intermediate filaments (NF200, green) and GABA (red). Scale bar, 200 µm. (B) The results of GABA^+^ and GABA-neurons quantification (n≥15 ROIs per condition, results obtained from 5 independent experiments) are expressed as mean±s.e.m.

**Figure S3.**
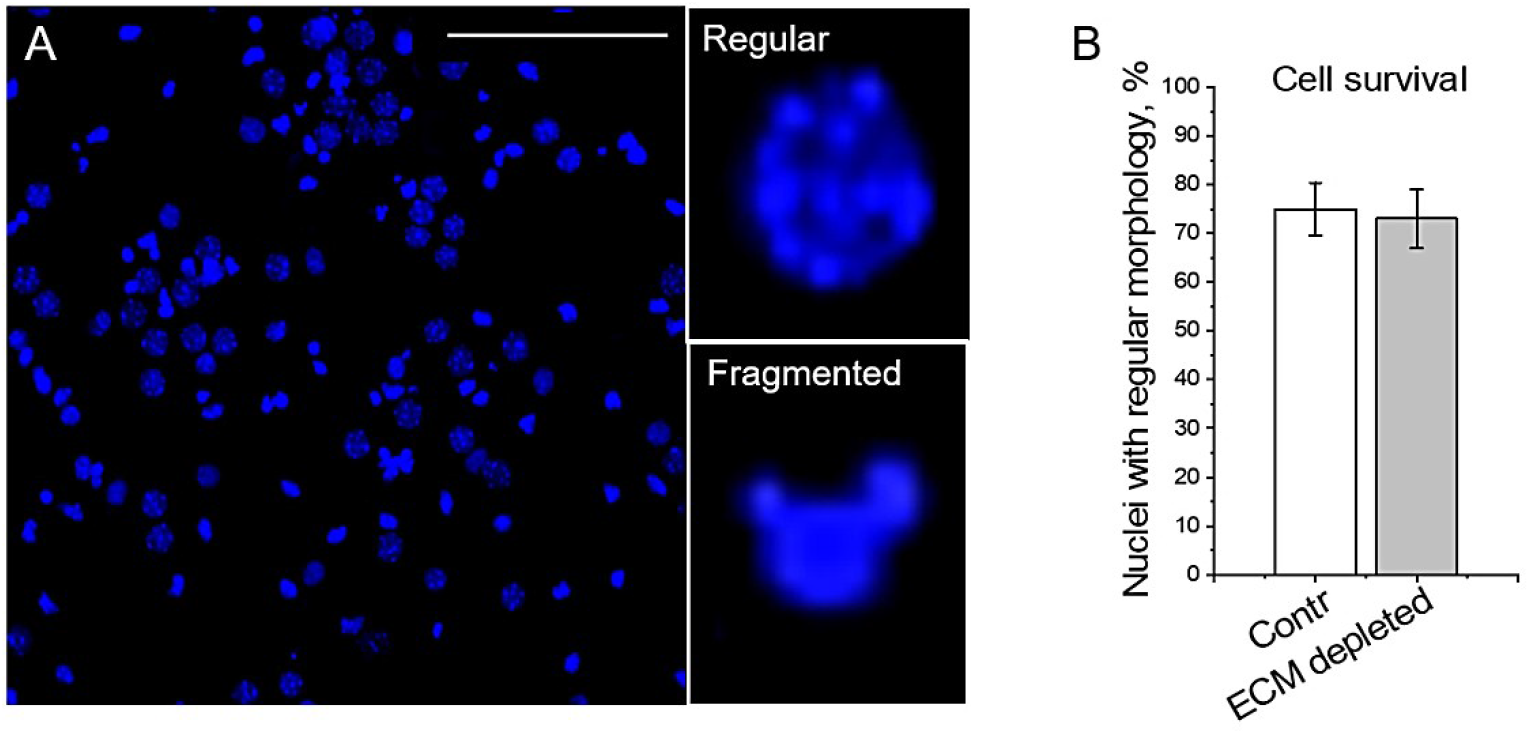
ECM depletion does not affect cell survival in neuronal cultures. (A) The micrograph shows representative labelling of nuclei (DAPI, blue). Examples of regular and fragmented nuclei are magnified. (B) A quantitative analysis or regular nuclei (n≥9 ROIs per condition, results obtained from 3 independent experiments) are expressed as mean±s.e.m.

**Figure S4.**
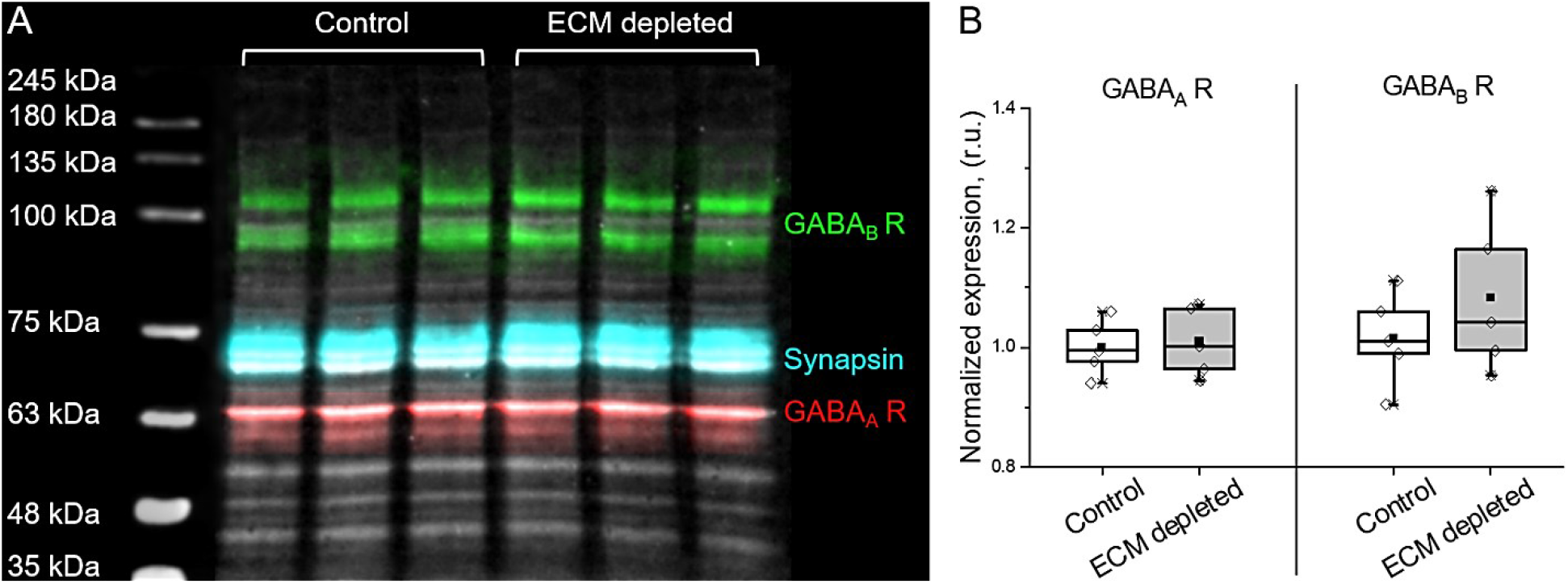
Cell surface expression of GABA receptors. (A) Multicolor detection of GABA_B_ receptors (green), GABA_A_ receptors (red), synapsin (cyan) and stain-free total protein (grey) on a single Western blot membrane, on which protein samples from cell membranes were loaded. (B) The total relative expression of surface GABA receptors is not altered by ECM depletion (n=5 independent Western blots).

**Figure S5.**
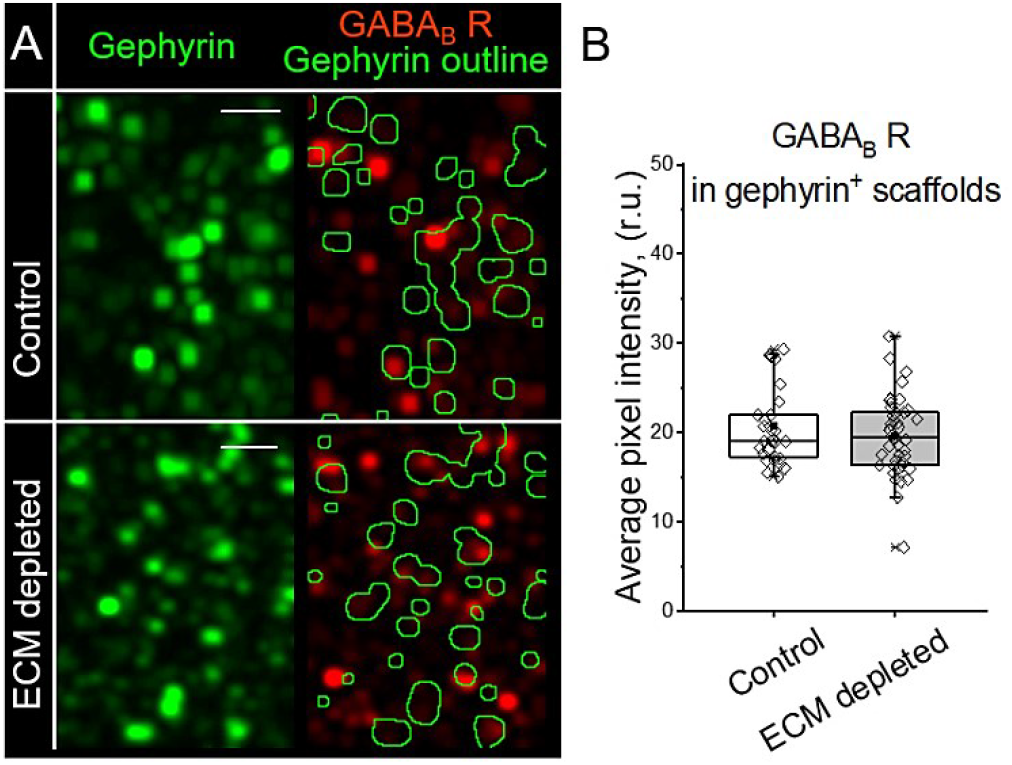
Postsynaptic expression of GABA_B_ receptors. (A) The panel illustrates the analysis of GABA_B_ receptor expression in inhibitory postsynapses (that is, gephyrin^+^ areas). The outlined areas (green) depict the regions in which the immunoreactivity of GABA_B_ receptors was measured. Scale bars, 2 μm. (B) ECM depletion does not alter the postsynaptic of GABA_B_ receptors. The average pixel intensity was quantified for each neuron examined (n≥ 30 cells per condition, results obtained from 5 independent experiments). Data are medians (lines inside boxes)/ means (filled squares inside boxes) ± IQR (boxes) with 10/ 90% ranks as whiskers. Open diamonds are data points.

**Figure S6.**
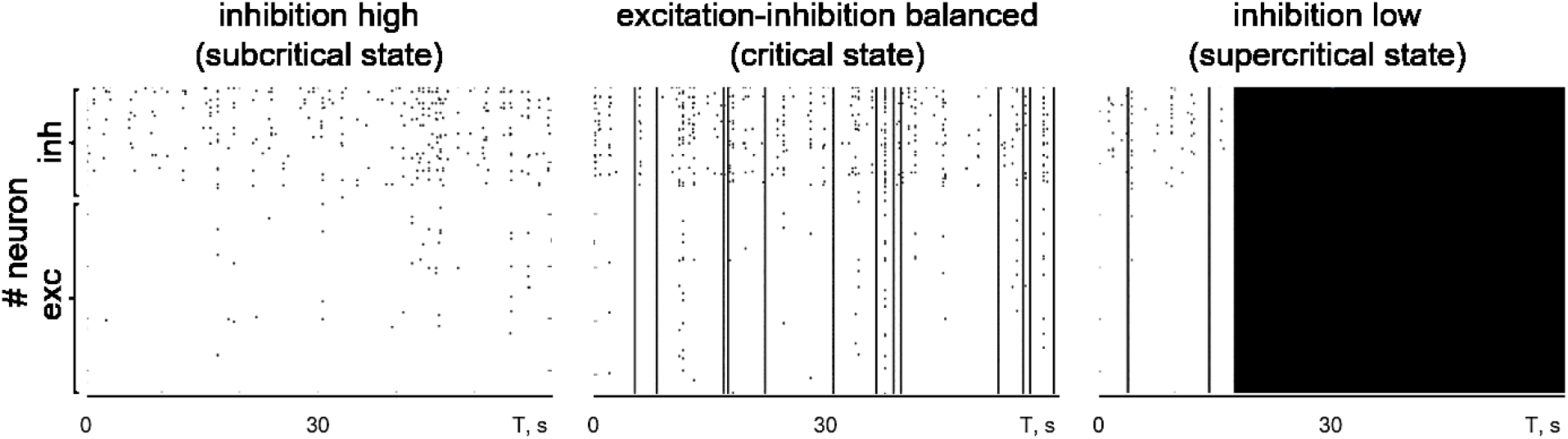
Criticality of in silico neuronal networks. The balance between excitatory and inhibitory inputs results in the stable spiking-bursting transitions characteristic for the critical state of the network. The disruption of E-I balance can result in network silencing (subcritical state) or uncontrolled synchronous firing (supercritical state). exc, excitatory; inh, inhibitory.

**Figure S7.**
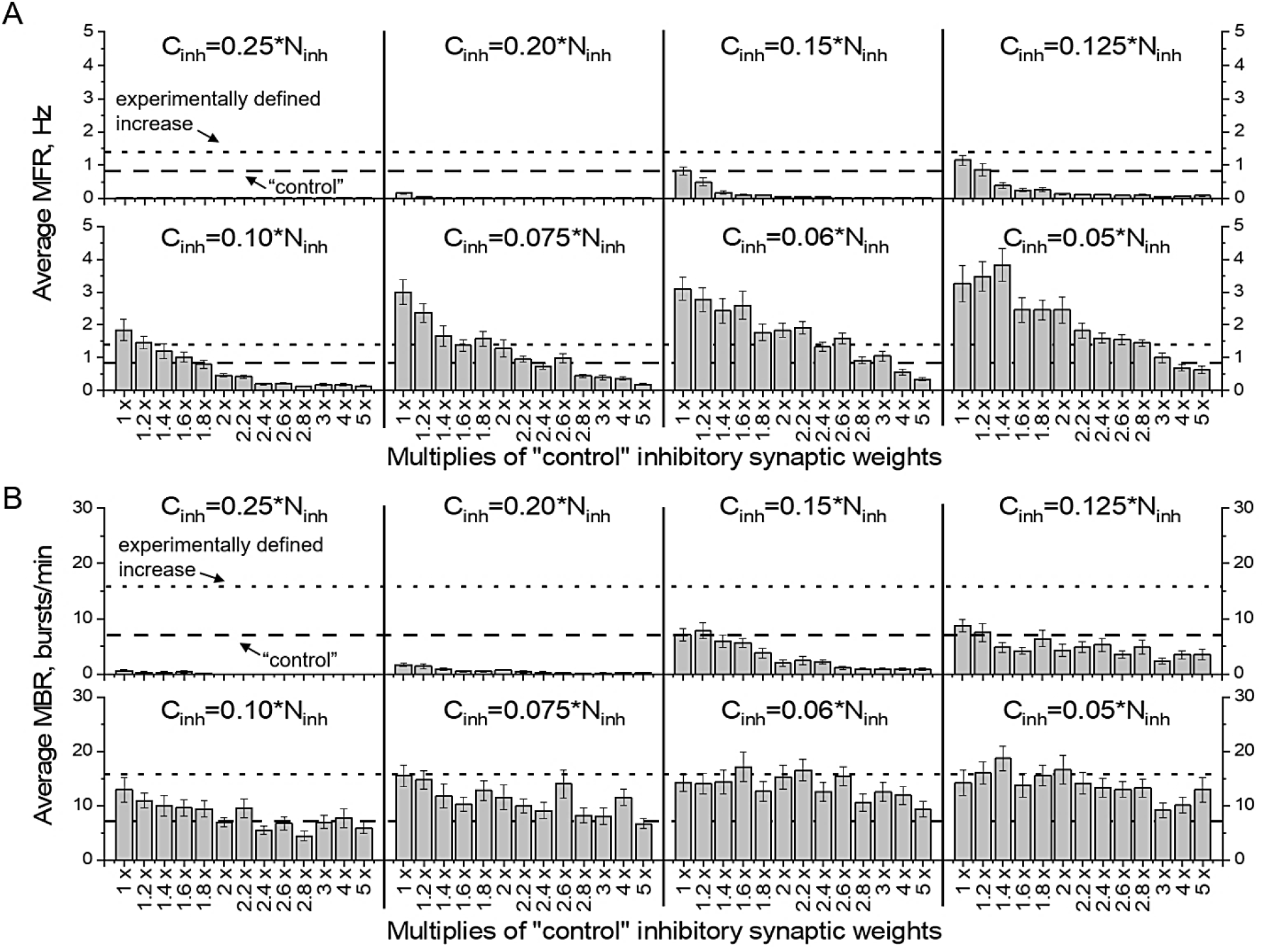
Impact of inhibitory connectivity and synapse strength changes on network activity. Quantifications of average (A) MFR and (B) MBR are shown for a range of C_inh_ and W_inh_ parameters. The dashed line indicates the average value in “control” simulations, the dotted line shows the experimentally observed increase of network activity after ECM depletion *in vitro*. The bars are mean±s.e.m. For each condition, n=15 independent simulation experiments were performed.

**Figure S8.**
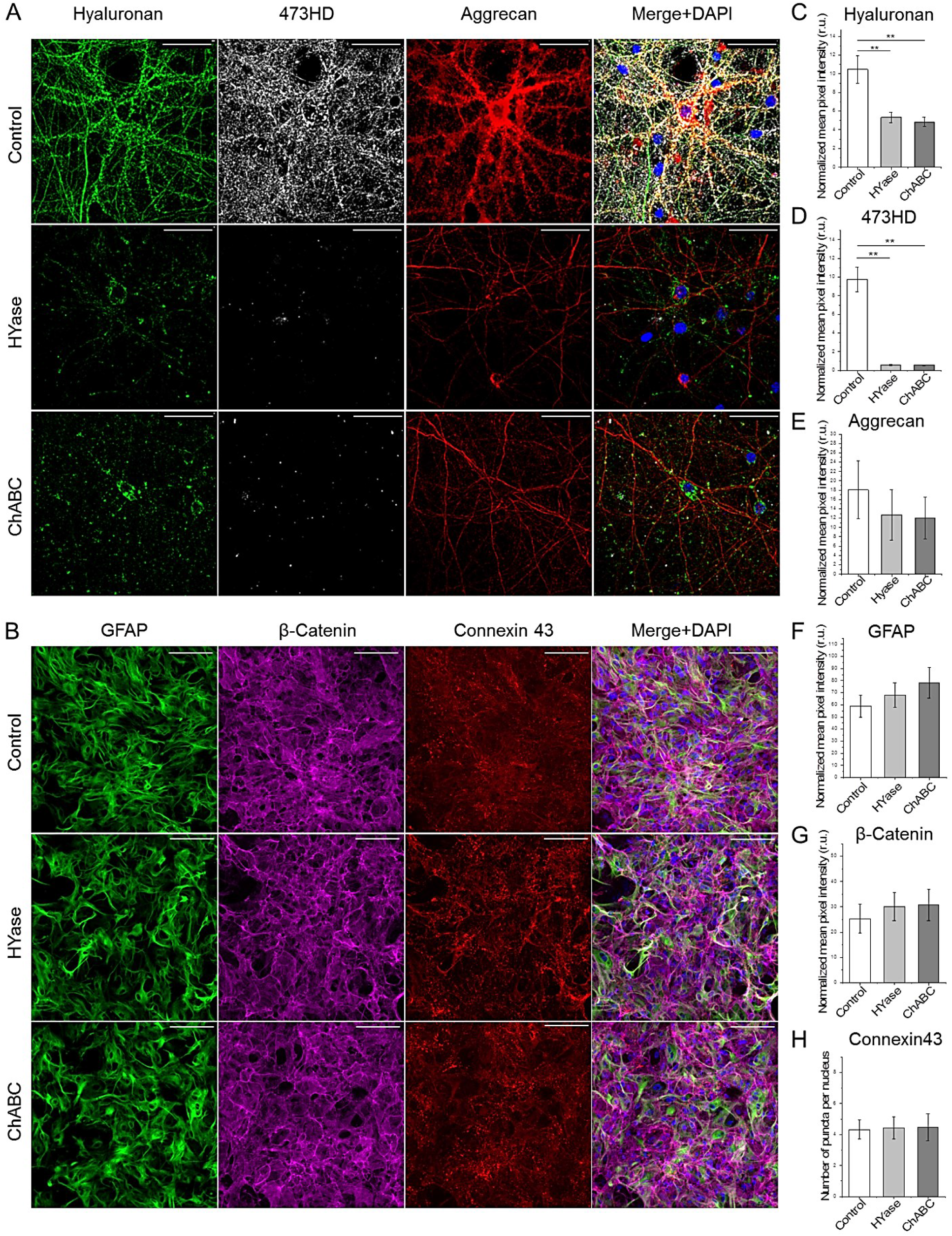
Enzymatic digestion removes neuronal ECM, but does not alter astrocyte morphology. (A) Representative immunostainings of hyaluronan, 473HD chondroitin sulfate epitope and aggrecan are shown under control condition and after treatment with hyaluronidase (Hyase) and chondroitinase ABC (ChABC). Scale bars, 50 µm. Both enzymatic treatments were equally efficient in depleting ECM components, as indicated by the quantifications of hyaluronan (C), 473HD chondroitin sulfate epitope (D) and aggrecan (E) labeling intensity. The results of quantification (n≥15 ROIs per condition, results obtained from 5 independent experiments) are expressed as mean±s.e.m. Asterisks indicate statistically significant differences, based on Kruskal-Wallis tests (**p<0.01). (B) Representative immunostainings of astrocytic monolayers expressing glial fibrillary acidic protein (GFAP), β-catenin and connexin 43 are shown under control condition and after treatment with Hyase and ChABC. Scale bars, 100 µm. The quantifications indicate that the expression of GFAP (F), β-catenin (G) and connexin 43 (H) was not altered after ECM digestion. The results of quantification (minimum 15 ROIs per condition, n=5) are expressed as mean±s.e.m.

**Figure S9.**
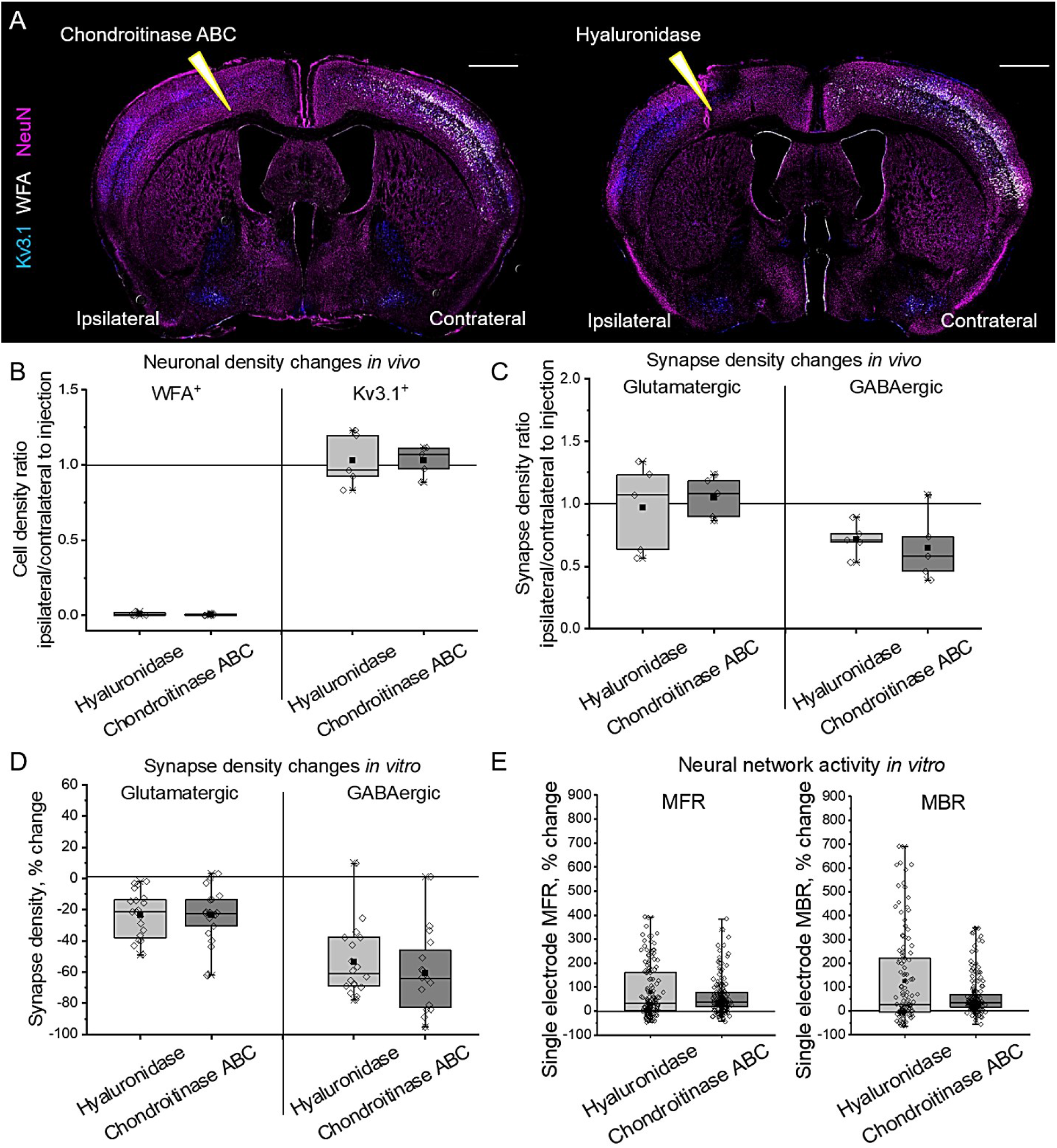
Enzymatic depletion of ECM with chondroitinase ABC or hyaluronidase induces similar synapse density and network activity alterations. (A) Representative immunolabelling of neuronal nuclei (NeuN, magenta), fast spiking interneurons (Kv3.1, blue) and PNNs (WFA, *Wisteria floribunda* agglutinin, white) is shown. Sharp triangles indicate intracortical injection sites. Squares indicate the regions in which cell and synapse densities were analyzed. Scale bars, 1 mm. (B) The loss of PNN expression and fast spiking interneurons was examined *in vivo*. Changes in PNN^+^ and Kv3.1^+^ neuron densities are expressed as ipsilateral to contralateral ratios. (C) Synapse density alterations *in vivo*. Changes in glutamatergic and GABAergic synapse densities are expressed as ipsilateral to contralateral ratios. Ratios in (B) and (C) are shown for each animal examined (n=5 animals per condition). (D) Synapse density alterations *in vitro*. Changes in glutamatergic and GABAergic synapse densities are expressed as differences with mean values of corresponding control experiments. Differences are shown for each neuron examined (n≥20 cells per condition, results obtained from 5 independent experiments). (E) Neural network activity changes *in vitro*. Mean firing rate (MFR) and mean bursting rate (MBR) changes are shown for single electrodes as differences with baseline activities of the same electrodes before treatment (n≥169 electrodes per condition, results obtained from 5 independent experiments). Data are medians (lines inside boxes)/ means (filled squares inside boxes) ± IQR (boxes) with 10/ 90% ranks as whiskers. Open diamonds are data points. No significant differences were detected, based on Kruskal-Wallis tests.

## Supplementary Code 1

The following code is a ready-to use MatLab script for performing network activity simulations

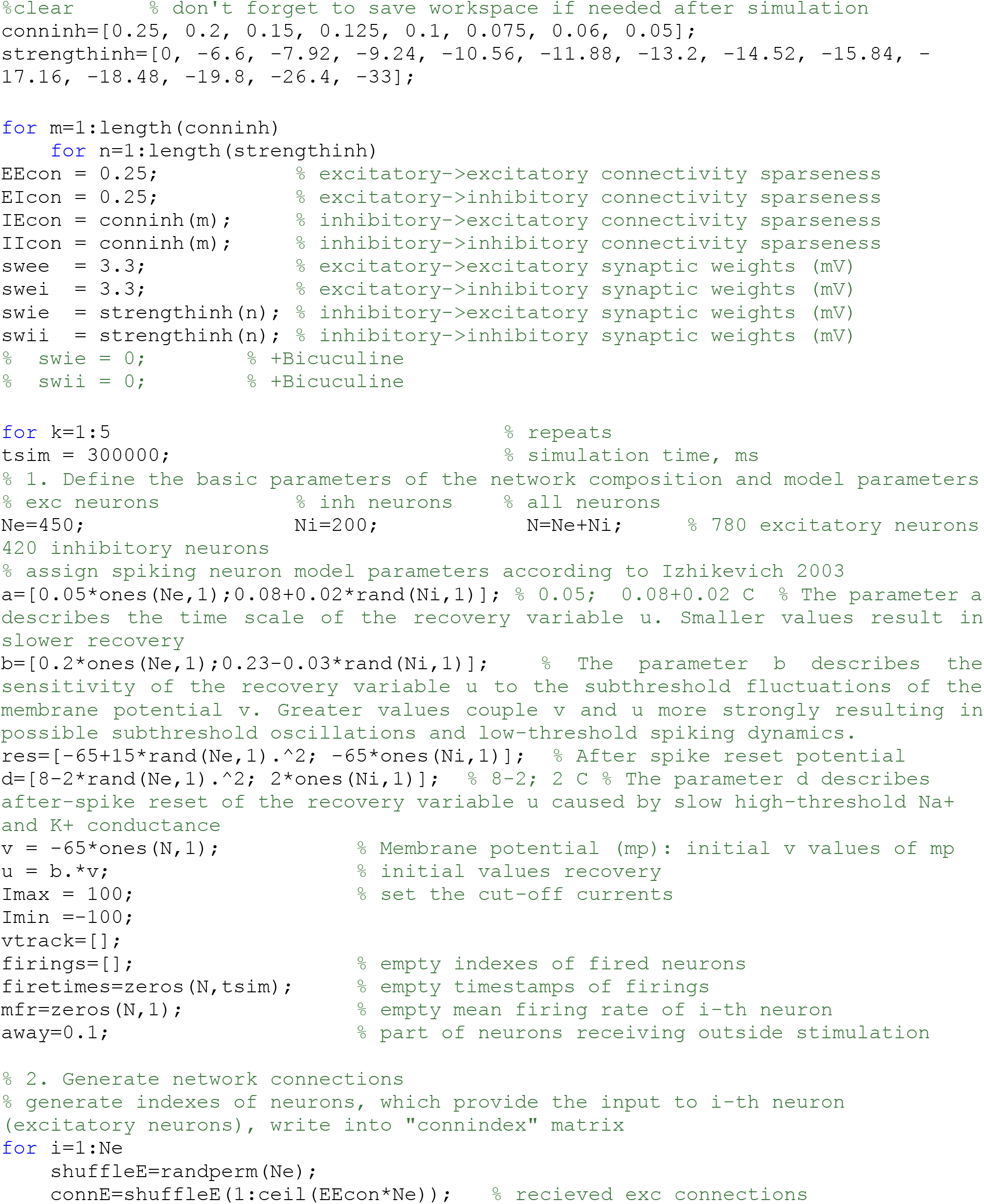
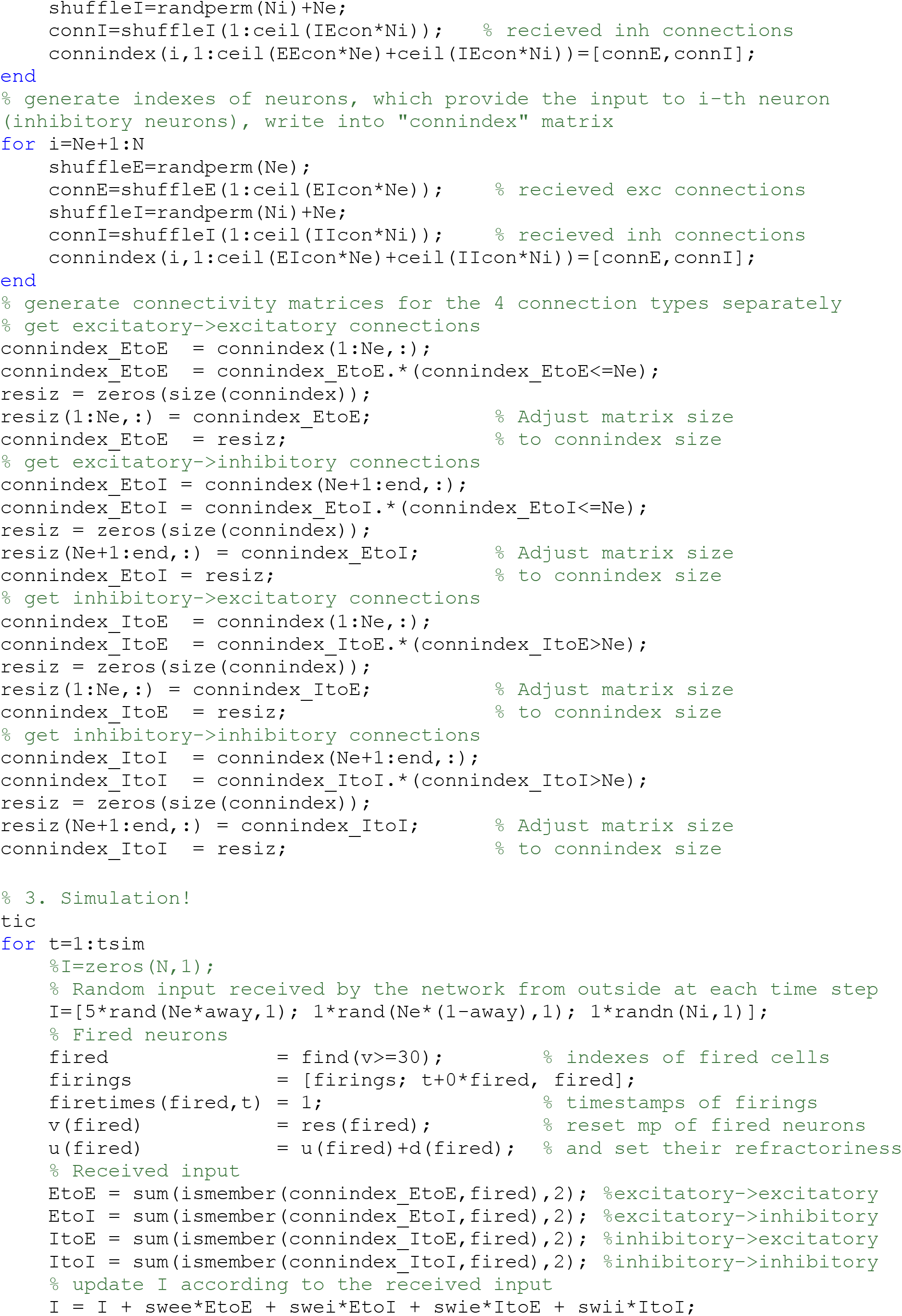
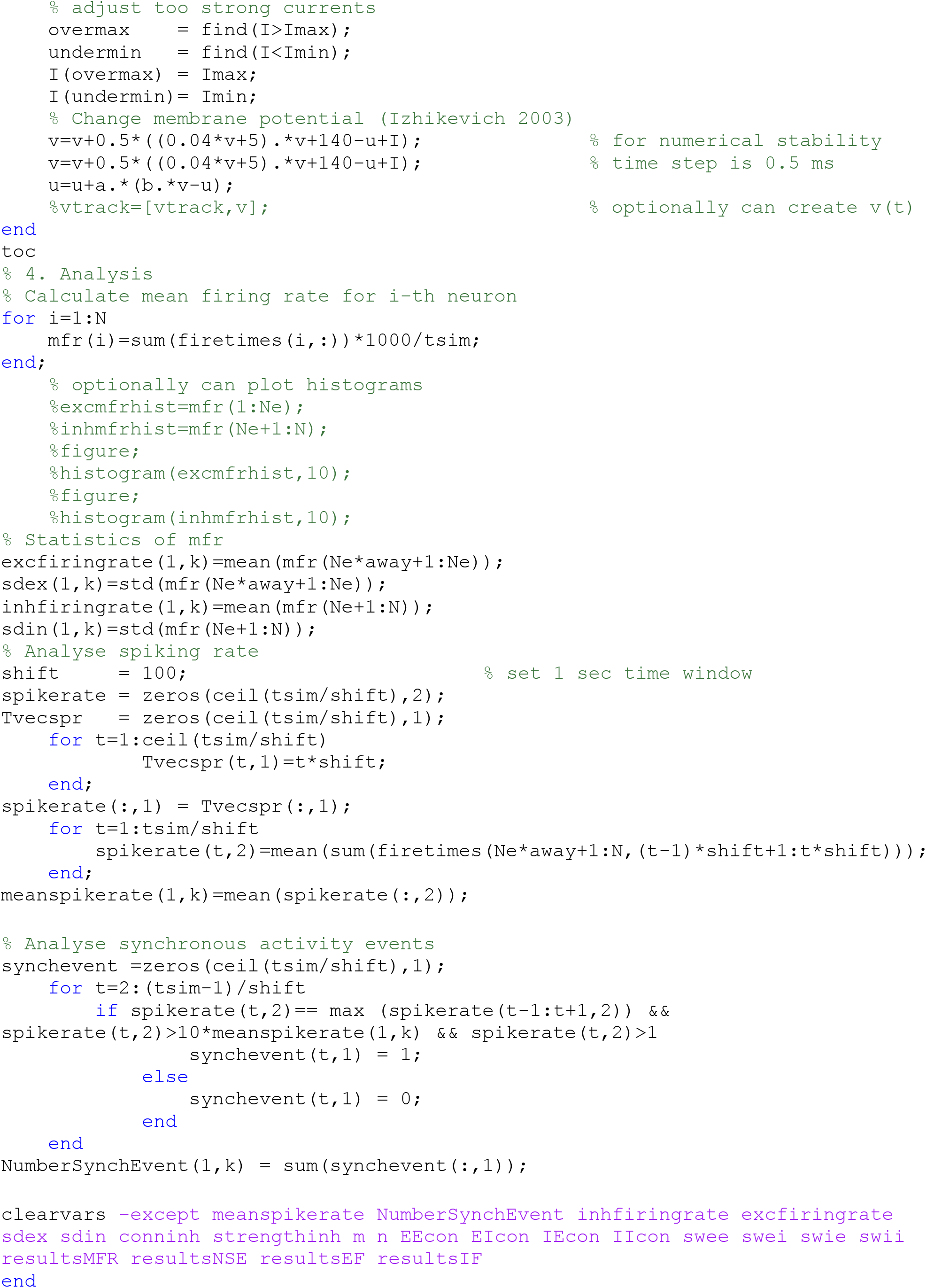
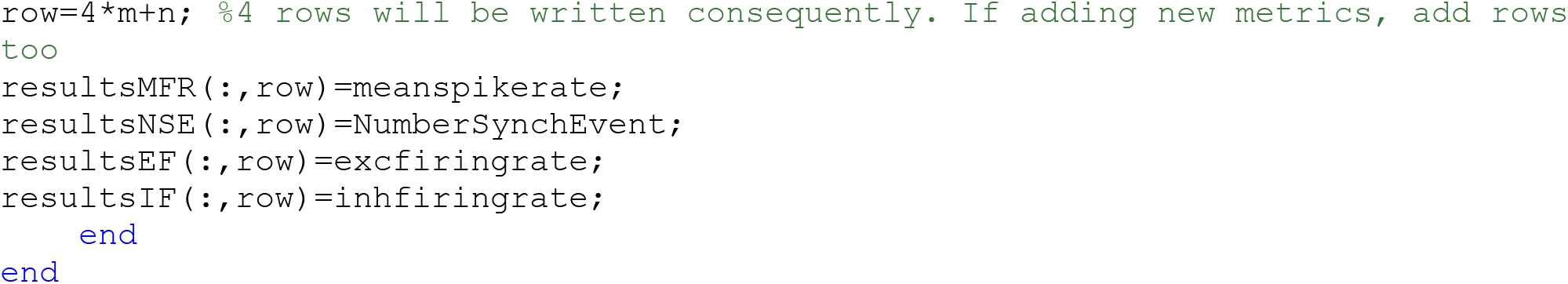

## Citations

Arranz, A.M., K.L. Perkins, F. Irie, D.P. Lewis, J. Hrabe, F. Xiao, N. Itano, K. Kimata, S. Hrabetova, and Y. Yamaguchi. 2014. Hyaluronan deficiency due to Has3 knock-out causes altered neuronal activity and seizures via reduction in brain extracellular space. The Journal of neuroscience : the official journal of the Society for Neuroscience. 34:6164–6176.

Bhatia, A., S. Moza, and U.S. Bhalla. 2019. Precise excitation-inhibition balance controls gain and timing in the hippocampus. eLife. 8:e43415.

Bikbaev, A., R. Frischknecht, and M. Heine. 2015. Brain extracellular matrix retains connectivity in neuronal networks. Sci Rep. 5:14527.

Blein, S., R. Ginham, D. Uhrin, B.O. Smith, D.C. Soares, S. Veltel, R.A. McIlhinney, J.H. White, and P.N. Barlow. 2004. Structural analysis of the complement control protein (CCP) modules of GABA(B) receptor 1a: only one of the two CCP modules is compactly folded. J Biol Chem. 279:48292–48306.

Bologna, L.L., V. Pasquale, M. Garofalo, M. Gandolfo, P.L. Baljon, A. Maccione, S. Martinoia, and M. Chiappalone. 2010. Investigating neuronal activity by SPYCODE multi-channel data analyzer. Neural Networks. 23:685–697.

Carulli, D., R. Broersen, F. de Winter, E.M. Muir, M. Meskovic, M. de Waal, S. de Vries, H.J. Boele, C.B. Canto, C.I. De Zeeuw, and J. Verhaagen. 2020. Cerebellar plasticity and associative memories are controlled by perineuronal nets. Proc Natl Acad Sci U S A. 117:6855–6865.

Carulli, D., K.E. Rhodes, D.J. Brown, T.P. Bonnert, S.J. Pollack, K. Oliver, P. Strata, and J.W. Fawcett. 2006. Composition of perineuronal nets in the adult rat cerebellum and the cellular origin of their components. J Comp Neurol. 494:559–577.

Carulli, D., K.E. Rhodes, and J.W. Fawcett. 2007. Upregulation of aggrecan, link protein 1, and hyaluronan synthases during formation of perineuronal nets in the rat cerebellum. Journal of Comparative Neurology. 501:83–94.

Choi, S.-Y. 2018. Synaptic and circuit development of the primary sensory cortex. Experimental & Molecular Medicine. 50:13.

Choii, G., and J. Ko. 2015. Gephyrin: a central GABAergic synapse organizer. Exp Mol Med. 47:e158.

Chu, P., R. Abraham, K. Budhu, U. Khan, N. De Marco Garcia, and J.C. Brumberg. 2018. The Impact of Perineuronal Net Digestion Using Chondroitinase ABC on the Intrinsic Physiology of Cortical Neurons. Neuroscience. 388:23–35.

Clarkson, A.N., B.S. Huang, S.E. Macisaac, I. Mody, and S.T. Carmichael. 2010. Reducing excessive GABA-mediated tonic inhibition promotes functional recovery after stroke. Nature. 468:305–309.

Davies, C.H., S.N. Davies, and G.L. Collingridge. 1990. Paired-pulse depression of monosynaptic GABA-mediated inhibitory postsynaptic responses in rat hippocampus. J Physiol. 424:513–531.

Dityatev, A., G. Bruckner, G. Dityateva, J. Grosche, R. Kleene, and M. Schachner. 2007. Activity-dependent formation and functions of chondroitin sulfate-rich extracellular matrix of perineuronal nets. Developmental neurobiology. 67:570–588.

Dzyubenko, E., C. Gottschling, and A. Faissner. 2016a. Neuron-Glia Interactions in Neural Plasticity: Contributions of Neural Extracellular Matrix and Perineuronal Nets. Neural Plast. 2016:5214961.

Dzyubenko, E., G. Juckel, and A. Faissner. 2017. The antipsychotic drugs olanzapine and haloperidol modify network connectivity and spontaneous activity of neural networks in vitro. Sci Rep. 7:11609.

Dzyubenko, E., D. Manrique-Castano, C. Kleinschnitz, A. Faissner, and D.M. Hermann. 2018. Topological remodeling of cortical perineuronal nets in focal cerebral ischemia and mild hypoperfusion. Matrix Biol. 74:121–132.

Dzyubenko, E., A. Rozenberg, D.M. Hermann, and A. Faissner. 2016b. Colocalization of synapse marker proteins evaluated by STED-microscopy reveals patterns of neuronal synapse distribution in vitro. J Neurosci Methods. 273:149–159.

Faini, G., A. Aguirre, S. Landi, D. Lamers, T. Pizzorusso, G.M. Ratto, C. Deleuze, and A. Bacci. 2018. Perineuronal nets control visual input via thalamic recruitment of cortical PV interneurons. eLife. 7:e41520.

Fawcett, J.W., T. Oohashi, and T. Pizzorusso. 2019. The roles of perineuronal nets and the perinodal extracellular matrix in neuronal function. Nat Rev Neurosci. 20:451–465.

Frerking, M., S. Borges, and M. Wilson. 1995. Variation in GABA mini amplitude is the consequence of variation in transmitter concentration. Neuron. 15:885–895.

Frischknecht, R., M. Heine, D. Perrais, C.I. Seidenbecher, D. Choquet, and E.D. Gundelfinger. 2009. Brain extracellular matrix affects AMPA receptor lateral mobility and short-term synaptic plasticity. Nat Neurosci. 12:897–904.

Gandolfi, D., A. Bigiani, C.A. Porro, and J. Mapelli. 2020. Inhibitory Plasticity: From Molecules to Computation and Beyond. Int J Mol Sci. 21.

Gautam, S.H., T.T. Hoang, K. McClanahan, S.K. Grady, and W.L. Shew. 2015. Maximizing Sensory Dynamic Range by Tuning the Cortical State to Criticality. PLoS Comput Biol. 11:e1004576.

Gottschling, C., E. Dzyubenko, M. Geissler, and A. Faissner. 2016. The Indirect Neuron-astrocyte Coculture Assay: An In Vitro Set-up for the Detailed Investigation of Neuron-glia Interactions. J Vis Exp:e54757.

Gottschling, C., D. Wegrzyn, B. Denecke, and A. Faissner. 2019. Elimination of the four extracellular matrix molecules tenascin-C, tenascin-R, brevican and neurocan alters the ratio of excitatory and inhibitory synapses. Sci Rep. 9:13939.

Haider, B., A. Duque, A.R. Hasenstaub, and D.A. McCormick. 2006. Neocortical network activity in vivo is generated through a dynamic balance of excitation and inhibition. The Journal of neuroscience : the official journal of the Society for Neuroscience. 26:4535–4545.

Härtig, W., B. Mages, S. Aleithe, B. Nitzsche, S. Altmann, H. Barthel, M. Krueger, and D. Michalski. 2017. Damaged Neocortical Perineuronal Nets Due to Experimental Focal Cerebral Ischemia in Mice, Rats and Sheep. Frontiers in Integrative Neuroscience. 11.

Ho, V.M., J.-A. Lee, and K.C. Martin. 2011. The Cell Biology of Synaptic Plasticity. Science. 334:623–628.

Hylin, M.J., S.A. Orsi, A.N. Moore, and P.K. Dash. 2013. Disruption of the perineuronal net in the hippocampus or medial prefrontal cortex impairs fear conditioning. Learn Mem. 20:267–273.

Izhikevich, E.M. 2003. Simple model of spiking neurons. IEEE Trans Neural Netw. 14:1569–1572.

Kinouchi, O., and M. Copelli. 2006. Optimal dynamical range of excitable networks at criticality. Nature physics. 2:348–351.

Krishnaswamy, V.R., A. Benbenishty, P. Blinder, and I. Sagi. 2019. Demystifying the extracellular matrix and its proteolytic remodeling in the brain: structural and functional insights. Cellular and Molecular Life Sciences. 76:3229–3248.

Lake, E.M.R., P. Bazzigaluppi, J. Mester, L.A.M. Thomason, R. Janik, M. Brown, J. McLaurin, P.L. Carlen, D. Corbett, G.J. Stanisz, and B. Stefanovic. 2017. Neurovascular unit remodelling in the subacute stage of stroke recovery. Neuroimage. 146:869–882.

Ma, Z., G.G. Turrigiano, R. Wessel, and K.B. Hengen. 2019. Cortical Circuit Dynamics Are Homeostatically Tuned to Criticality In Vivo. Neuron. 104:655.664 e654.

Mongillo, G., S. Rumpel, and Y. Loewenstein. 2018. Inhibitory connectivity defines the realm of excitatory plasticity. Nat Neurosci. 21:1463–1470.

Nahmani, M., and A. Erisir. 2005. VGluT2 immunochemistry identifies thalamocortical terminals in layer 4 of adult and developing visual cortex. J Comp Neurol. 484:458–473.

Nusser, Z., S. Cull-Candy, and M. Farrant. 1997. Differences in Synaptic GABAA Receptor Number Underlie Variation in GABA Mini Amplitude. Neuron. 19:697–709.

Pantazopoulos, H., M. Markota, F. Jaquet, D. Ghosh, A. Wallin, A. Santos, B. Caterson, and S. Berretta. 2015. Aggrecan and chondroitin-6-sulfate abnormalities in schizophrenia and bipolar disorder: a postmortem study on the amygdala. Transl Psychiatry. 5:e496.

Pastore, V.P., P. Massobrio, A. Godjoski, and S. Martinoia. 2018. Identification of excitatory-inhibitory links and network topology in large-scale neuronal assemblies from multi-electrode recordings. PLOS Computational Biology. 14:e1006381.

Pfeiffer, T., S. Poll, S. Bancelin, J. Angibaud, V.K. Inavalli, K. Keppler, M. Mittag, M. Fuhrmann, and U.V. Nagerl. 2018. Chronic 2P-STED imaging reveals high turnover of dendritic spines in the hippocampus in vivo. Elife. 7.

Pitler, T.A., and B.E. Alger. 1994. Differences between presynaptic and postsynaptic GABAB mechanisms in rat hippocampal pyramidal cells. J Physiol. 72:2317–2327.

Pizzorusso, T., P. Medini, N. Berardi, S. Chierzi, J.W. Fawcett, and L. Maffei. 2002. Reactivation of ocular dominance plasticity in the adult visual cortex. Science. 298:1248–1251.

Pless, E. 2009. An investigation of interactions with extracellular matrix proteins mediated by the CCP modules of the metabotropic GABAB receptor.

Roberto, M., S.G. Madamba, D.G. Stouffer, L.H. Parsons, and G.R. Siggins. 2004. Increased GABA release in the central amygdala of ethanol-dependent rats. The Journal of neuroscience : the official journal of the Society for Neuroscience. 24:10159–10166.

Roll, L., and A. Faissner. 2014. Influence of the extracellular matrix on endogenous and transplanted stem cells after brain damage. Frontiers in Cellular Neuroscience. 8.

Shi, W., X. Wei, X. Wang, S. Du, W. Liu, J. Song, and Y. Wang. 2019. Perineuronal nets protect long-term memory by limiting activity-dependent inhibition from parvalbumin interneurons. Proceedings of the National Academy of Sciences. 116:27063–27073.

Sigal, Y.M., H. Bae, L.J. Bogart, T.K. Hensch, and X. Zhuang. 2019. Structural maturation of cortical perineuronal nets and their perforating synapses revealed by superresolution imaging. Proceedings of the National Academy of Sciences. 116:7071–7076.

Soleman, S., M.A. Filippov, A. Dityatev, and J.W. Fawcett. 2013. Targeting the neural extracellular matrix in neurological disorders. Neuroscience. 253:194–213.

Sprekeler, H. 2017. Functional consequences of inhibitory plasticity: homeostasis, the excitation-inhibition balance and beyond. Current opinion in neurobiology. 43:198–203.

Tewari, B.P., L. Chaunsali, S.L. Campbell, D.C. Patel, A.E. Goode, and H. Sontheimer. 2018. Perineuronal nets decrease membrane capacitance of peritumoral fast spiking interneurons in a model of epilepsy. Nat Commun. 9:4724.

Trapp, P., R. Echeveste, and C. Gros. 2018. E-I balance emerges naturally from continuous Hebbian learning in autonomous neural networks. Sci Rep. 8:8939.

Villa, K.L., K.P. Berry, J. Subramanian, J.W. Cha, W.C. Oh, H.B. Kwon, Y. Kubota, P.T. So, and E. Nedivi. 2016. Inhibitory Synapses Are Repeatedly Assembled and Removed at Persistent Sites In Vivo. Neuron. 89:756–769.

von Holst, A., S. Sirko, and A. Faissner. 2006. The unique 473HD-Chondroitinsulfate epitope is expressed by radial glia and involved in neural precursor cell proliferation. The Journal of neuroscience : the official journal of the Society for Neuroscience. 26:4082–4094.

Wang, Y.C., E. Dzyubenko, E.H. Sanchez-Mendoza, M. Sardari, T. Silva de Carvalho, T.R. Doeppner, B. Kaltwasser, P. Machado, C. Kleinschnitz, C.L. Bassetti, and D.M. Hermann. 2018. Postacute Delivery of GABAA alpha5 Antagonist Promotes Postischemic Neurological Recovery and Peri-infarct Brain Remodeling. Stroke. 49:2495–2503.

